# Cascades and convergence: dynamic signal flow in a synapse-level brain network

**DOI:** 10.1101/2024.12.04.624189

**Authors:** Richard F. Betzel, Maria Grazia Puxeddu, Caio Seguin, Bratislav Misic

## Abstract

Connectomes—maps of synaptic connectivity—constrain how signals flow through the nervous system, shaping the transmission of sensory information to downstream targets involved in perception, decision-making, and action. Here, we use a simple, network-based spreading model to simulate sensory signal propagation across the adult Drosophila connectome. This approach allows us to trace modality-specific cascades and quantify their zones of overlap—neurons activated by multiple sensory pathways. Extending the classical spreading model, we introduce cooperative and competitive dynamics to simulate multisensory integration scenarios. Finally, we classify neurons based on their dynamical response profiles across all simulations, yielding a data-driven taxonomy grounded in both structure and dynamics. Our results highlight how abstract models can reveal organizing principles of neural computation and generate hypotheses for future experimental validation.

## INTRODUCTION

Connectomes are wiring maps that describe the connectivity of neural elements—cells, populations, areas—to one another [1–6]. These physical connections support signaling and communication across neural systems [7–10], enabling the transformation of sensory inputs into adaptive behavioral responses [11–14]. Identifying the pathways that govern this information flow is a central aim of systems and behavioral neuroscience [15, 16].

Network neuroscience provides a framework for representing and analyzing brain networks using tools from graph theory [17–20]. Here, neural elements are encoded as nodes and their pairwise connections as edges in a directed, weighted graph. Such models can support both structural analyses and simulations of network-constrained dynamics [21–24]. One class of models, spreading dynamics, imagines that activation initiated in a small set of seed nodes propagates stepwise across synaptic connections, resulting in activation cascades that spread through the network [25–27]. These simplified models offer analytical tractability and insight into how network architecture shapes large-scale dynamics. They also provide an idealized description of sensory processing—tracking how input from peripheral sensory neurons flows through the brain to drive downstream activity [28, 29].

Recent advances in large-scale electron microscopy have yielded nanoscale connectomes of entire nervous systems [30–34]. The largest and most complete to date is that of the adult female fruit fly, *Drosophila melanogaster*, comprising roughly 138,000 neurons and 54 million synapses [35]. In addition to its scale, the connectome features detailed cell-type annotations and modular subdivisions [36– enabling integrative analyses that link structure, function, and lineage.

In *Drosophila*, sensory systems are initially segregated by modality, but their signals converge onto shared higher-order structures such as the lateral horn, mushroom body, and central complex [39– These regions are thought to support associative learning, multisensory integration, and behavioral decision-making. Behavioral and physiological studies have shown that flies integrate information across sensory channels [42], though the circuit-level mechanisms remain poorly understood.

Using anatomical annotations of the fly connectome, it is now possible to identify sensory neurons and track how activity initiated in specific modalities spreads through the brain. In this study, we use and extend a recently developed spreading model [43] to explore how signals propagate from distinct sensory systems. We examine three scenarios: (1) *unimodal cascades*, in which single modalities are seeded independently; (2) *cooperative cascades*, in which paired modalities jointly influence activation timing; and (3) *competitive cascades*, where pairs of modalities vie for influence over shared downstream targets. We show that individual sensory cascades follow distinct trajectories but converge in central structures, particularly the central complex. We identify the neurons and connections that facilitate early, rapid activation, and analyze which neurons lie at the interface between competing signals. Finally, we use cascade-based summary statistics to group neurons by their roles in signal propagation.

In summary, this study makes both descriptive and theoretical contributions. Descriptively, we document the spatiotemporal trajectories of sensory cascades and link them to specific annotations and network features. Our work also makes notable contributions to theory. We demonstrate that early propagation of sensory cascades are facilitated by high degree hubs, that sensory modalities can be grouped into families based on the spatiotemporal similarity of their sensory cascades, that neurons in the central brain serve as convergence zones for multi-modal sensory cascades and exhibit high degrees of entropy. More generally, our work underscores the critical role of the connectome alone, even in the absence of any exogenous control signal, for shaping the propagation of activity throughout a nervous system.

## RESULTS

Here we analyzed the connectome of a single adult female *Drosophila melanogaster* (publicly available through https://codex.flywire.ai/; all analyses were carried out on the v783 connectome) [35]. The connectome comprised *N* = 138, 639 neurons, *M* = 15, 091, 983 edges, *M*_*w*_ = 54, 492, 922 synapses. The connectome is, by construction, directed and its edges weighted by synapse count.

### Neuronal wiring of sensory systems

Synaptic connectivity between neurons enables the propagation of signals—including sensory information—throughout the nervous system. The configuration of these connections—the connectome’s topology—thus plays a central role in shaping information flow [44]. To investigate how network structure contributes to sensory integration, we first examine the wiring among *N*_*sensory*_ = 16, 349 annotated sensory neurons. These neurons span several distinct modalities: gustation, hygrosensation, mechanosensation, olfaction, thermosensation, vision (both optic lobe and ocellar), and a category labeled “unknown.”

We extracted the 16349 × 16349 subgraph comprising all synaptic connections between sensory neurons (Fig. 2a). This subgraph was notably sparse, with a binary connection density of *d*_*sensory*_ = 1.3 × 10^−4^—approximately six times lower than the full connectome’s density (*d*_*connectome*_ = 7.8 × 10^−4^). Moreover, it was significantly sparser than randomly sampled subgraphs of the same size (*p <* 10^−3^; 1000 random samples).

**Figure 1.**
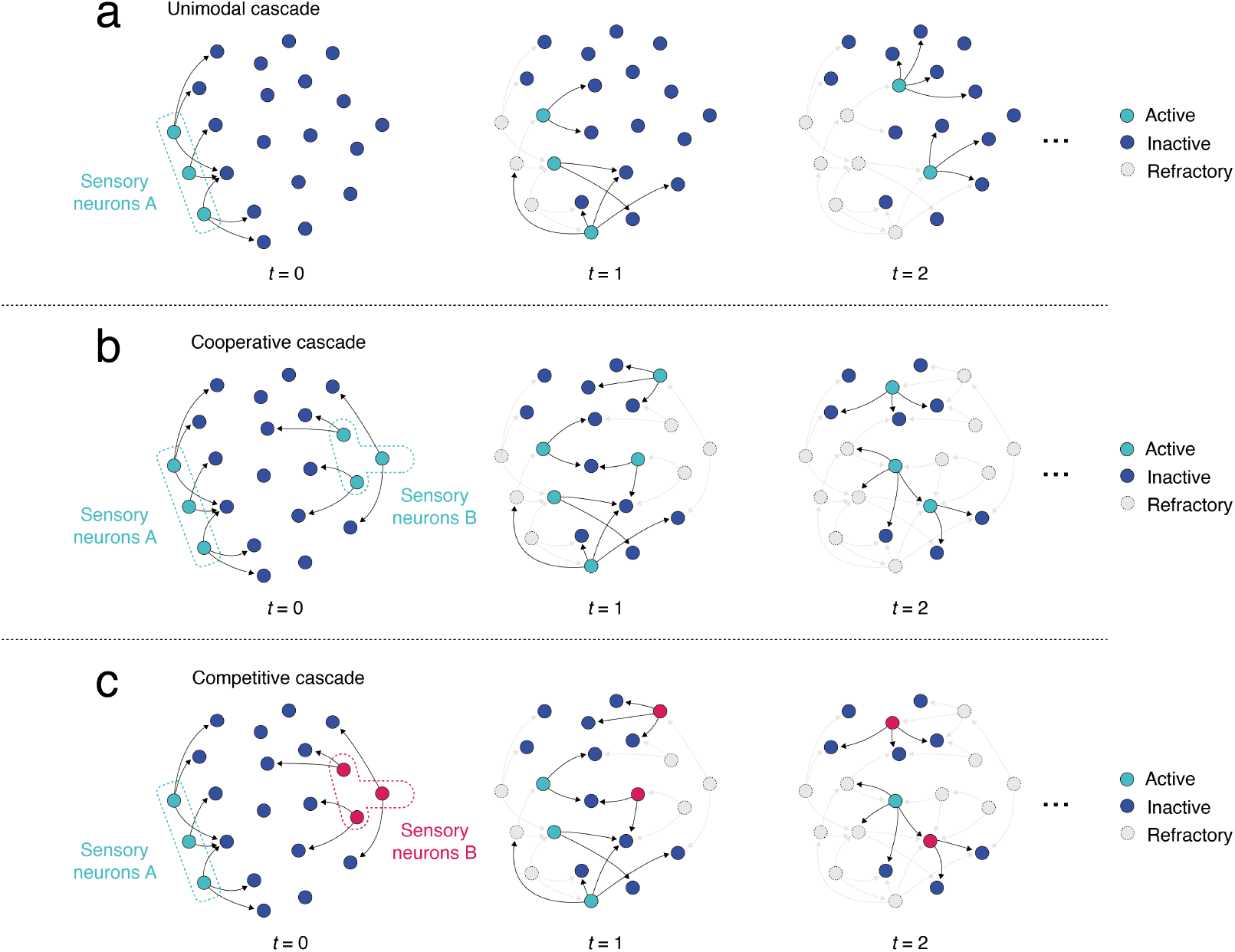
Schematic illustration of cascade model. Neurons can take on three states; active, inactive, or refractory. Inactive neurons are activated probabilistically by their pre-synaptic partners. Once a neuron has been activated, it enters a refractory period, wherein it cannot be activated at any subsequent time point. We consider three variations of this model. (*a*) Unimodal sensory cascades: wherein the initial population of active neurons (the seed population) are all of the same sensory modality. The model proceeds until either all neurons have been activated or there exists a time point wherein no neurons are currently active. (*b*) Cooperative sensory cascades: we consider two populations of sensory neurons, each of which represents a different sensory modality. Nonetheless, in their active states, these neurons carry identical information (the same state value), and therefore generally improve (reduce) activation times such that neuron *i* will tend to be activated earlier in the cooperative condition than in the unimodal condition. (*c*) Competitive sensory cascades: we, again, consider two populations of sensory neurons, but force them to carry different information and therefore “compete” for territory. Under the competitive model, it is possible for a neuron to be activated at the same time step by multiple neurons from both sensory cascades (green and red, in this panel). If this occurs, we count the number of green and red neurons. The neuron is assigned the color of the winning cascade. If there is an exact tie, then the winner is selected randomly.

**Figure 2.**
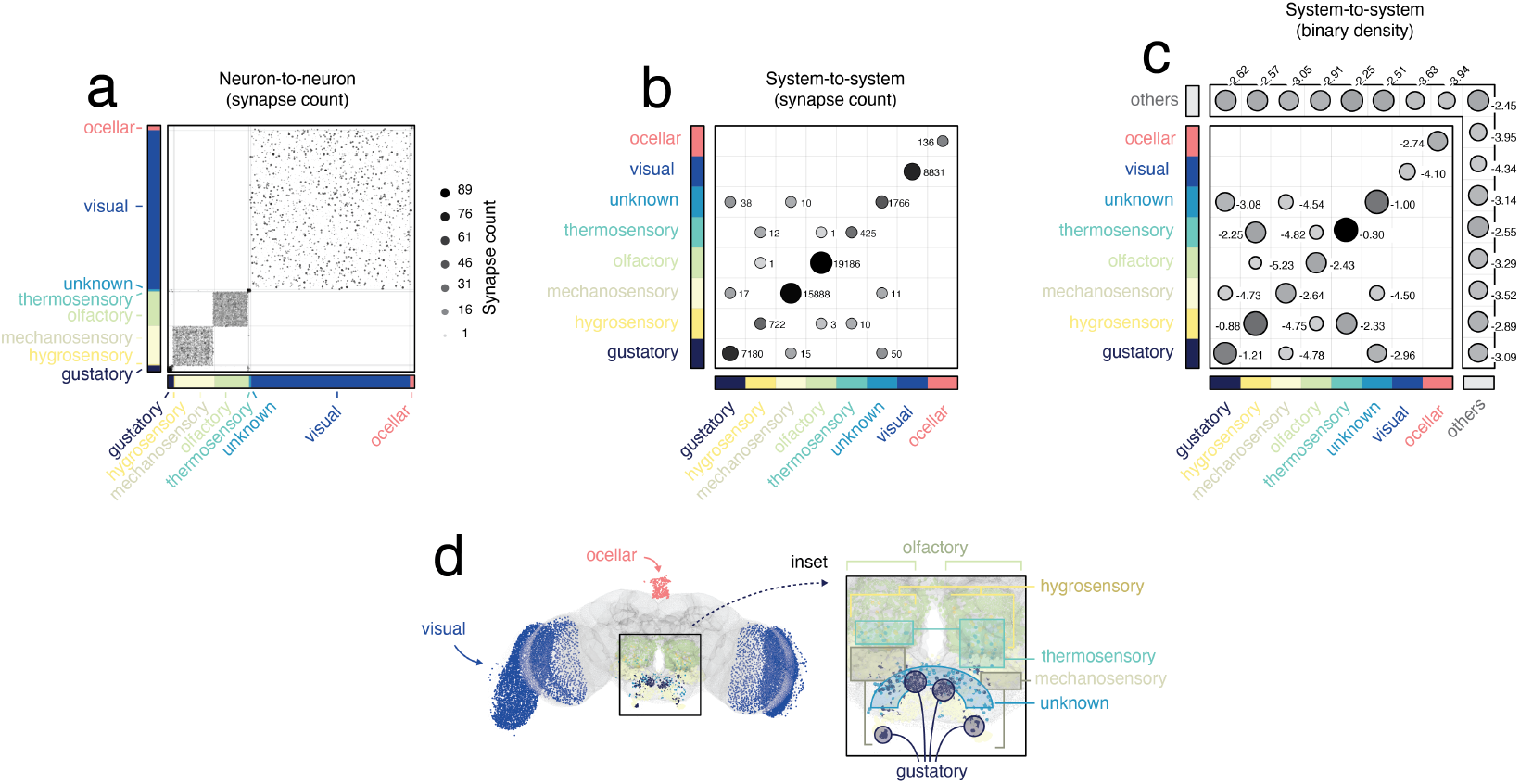
Synaptic connectivity within and between primary sensory neurons. (*a*) Complete sub-network of sensory-to-sensory connections. Gray lines delineate systems (modalities) from one another. (*b*) Synapse count between pairs of systems. (*c*) Binary density of connections between pairs of systems (and remaining neurons, labeled “others”). Note that in *c* the color and labels are on a logarithmic scale. (*d*) Anatomical depiction of sensory neurons.

The sensory subgraph also exhibited strong modularity. Of the 54,302 synapses, 54,134 (99.7%) connected neurons within the same modality (Fig. 2b). This high degree of within-modality wiring highlights a parallel, segregated architecture with minimal direct cross-talk between modalities.

Taken together, these results indicate that direct sensory-sensory interactions are both rare and strongly modality-specific. This organization suggests that multimodal integration likely occurs through polysynaptic pathways and that the secondary steps along these pathways involve non-sensory neurons.

To explore this possibility, we used a stochastic cascade model to simulate how activation might spread from sensory neurons across the full connectome. This abstract model is broadly applicable to networked systems [45], and provides a tractable framework for studying the spatiotemporal dynamics of signal propagation.

In the model, each neuron can be in one of three states: active, inactive, or refractory. At time *t* = 0, a small set of *N*_*seed*_ “seed” neurons is initialized as active—all others begin inactive. At time *t* = 1, the seed neurons can activate their post-synaptic partners. The *Drosophila* connectome is polyadic, meaning that the same pair of neurons can be connected by multiple synapses (at different locations along their processes). For pairs of neurons connected by more than one synapse, we assume that any of the synapses can activate the post-synaptic neuron. We treat each synapse as a Bernoulli trial where the probability of a “success” (activation of post-synaptic neuron) is equal to *p*_*transmission*_. Following activation, neurons turn off the next step (entering a “refractory” state), where they cannot activate their post-synaptic partners nor can they be activated by their pre-synaptic inputs at any point during the rest of the cascade. The cascade continues until no neurons are active. See Fig. S1 for example cascades and for details on the influence of model parameters. In the main text we fix parameter values to *p*_*transmission*_ = 0.01 and *N*_*seed*_ = 16. These particular values were selected practically; they are both large enough to ensure that the majority of cascades propagate to the entire network but still small enough to ensure that the cascades evolve over a long enough timescale to allow for meaningful analysis. We note, however, that alternative parameter values tend to stretch or contract activation timing, generally preserving rank order of activation (see Fig. S1d-f and Fig. S2).

### Edge contributions to cascades

The cascade model allows us to track how activity spreads not only across neurons, but also across individual synapses. This feature enables the identification of dynamic sensory pathways—sequences of synaptic contacts—that support the propagation of activation from sensory neurons into the broader brain. We can then ask: which types of neurons participate in these cascades? Do these pathways align with structural features of the connectome? And are the most active synapses associated with classical hub neurons?

To address these questions, we performed 1000 simulations of the cascade model in which *N*_*seed*_ = 16 sensory neurons were randomly drawn across all modalities and activated with *p*_*transmission*_ = 0.01. At each time step, we calculated across all simulations three quantities: the total number of active synapses, the number of unique synapses used, and their ratio (unique/total). Here, we consider a synapse to be active at time *t* if its post-synaptic neuron was inactive at *t* − 1 but activated at time *t*. At each time step, we measured three quantities: the total number of active synapses, the number of unique synapses used, and their ratio (unique/total). A high ratio would indicate consistent usage of the same synapses across simulations; a low ratio would suggest variable, diverse pathways. Note that while the seed neurons are drawn from all sensory modalities, we view this model as a special case of the unimodal model. The rationale for doing so is that, whereas the cooperative and competitive models explicitly test for synergistic effects of multi-sensory seeding and yield estimates of sensory “boundaries”, respectively, the summary statistics we would derive for this model–drawing seeds randomly from among all sensory modalities–are identical to the statistics used to summarize the unimodal model.

Early in the cascade, we observed a pronounced “bottlenecking” effect—specific synapses were repeatedly and consistently used to relay activity from the sensory seeds to other, secondary neurons (Fig. 3c). We show the most frequently used neurons and their connections at each of the first four steps in Fig. 3a and the average over all steps in Fig. 3b. This consistency declined at later stages, as the cascade diversified into multiple alternative routes (Fig. 3c). These findings suggest that a small set of high-traffic synapses mediates the initial wave of integration, while later dynamics engage a broader array of pathways.

**Figure 3.**
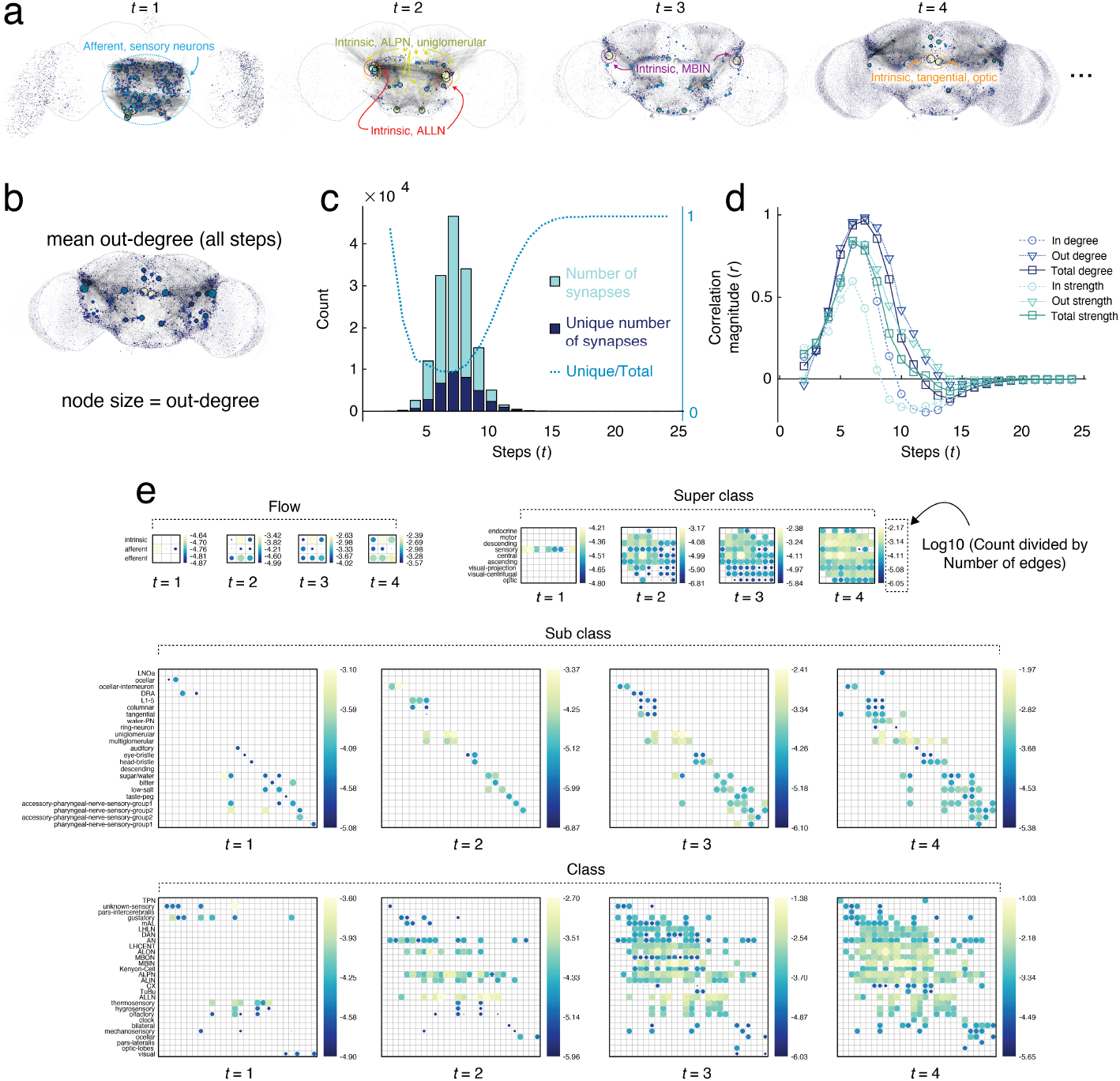
Edge sequencing. (*a*) Top 1000 edges (by usage) over first four time steps. (*b*) We constructed a weighted connectivity matrix by averaging edge usage across all time points, where at each time point an edge is present if it activates a postsynaptic neuron. We then calculate the out-degree of each node to highlight influential spreaders. (*c*) Across 1000 simulations, we count the total number of active synapses at each time point along with the unique number of active synapses. If the same synapses were active across all runs, then these numbers would coincide; however, due to the stochasticity of the cascade model and variation in the set of seeded neurons, different synapses appear at different time points. (*d*) Correlation of the instantaneous out-degree with basic structural properties of the network. (*e*) Active synapses (edges) grouped by annotation data. Circumference and color circle are proportional to the logarithm of their usage (count normalized by the number of synapses between pairs of annotations).

To examine whether synapse usage could be predicted from static network features, we computed the outdegree of each neuron based on edge usage (dynamic centrality) and compared this to their degree and strength in the underlying connectome (structural centrality). Early in the cascade, correlations between dynamic and structural centrality were weak (maximum *r* = 0.15), indicating that highly connected neurons in the static network did not dominate the initial spread (Fig. 3d). However, as activity progressed beyond *t* = 6, correlations rose sharply, exceeding *r >* 0.9 by *t* = 7, consistent with a transition toward greater reliance on structural hubs for sustaining propagation.

To better understand the cellular and anatomical substrates of these dynamics, we aggregated the 138,639 × 138,639 edge usage matrix at each time point by hierarchical annotation categories—”Flow”, “Superclass”, “Class”, and “Sub-class” (see **Materials and Methods**). Early cascade stages primarily involved synapses between afferent and intrinsic neurons, many of which were sensory (Fig. 3e, top panels). At intermediate time steps, uni- and multiglomerular projection neurons became prominent, including reciprocal interactions among these cell types and with tangential neurons—suggesting the involvement of antennal lobe circuits and early olfactory integration (Fig. 3e, middle). From *t* = 2 onward, activation increasingly involved downstream targets such as the mushroom body—particularly Kenyon cells—and neurons of the central complex. These populations serve as known convergence zones for multi-modal integration in *Drosophila* [46–48], and they relay activity to neurons in the optic lobes at later stages (Fig. 3e, bottom). These dynamics suggest a structured flow from afferent sensory neurons through intermediate integration zones to-ward broader, distributed activation of the brain.

### Overlapping and distinct spatiotemporal profiles of unimodal sensory cascades

In the previous section, we examined cascade dynamics following simultaneous activation of neurons from *all* sensory systems. Here, we restrict seeding to individual modalities, allowing us to isolate features that are shared across, or unique to, specific sensory systems.

To illustrate modality-specific dynamics, we compare cascades seeded from gustatory and hygrosensory neurons by visualizing their activation probability maps from *t* = 2 to *t* = 6 (Fig. 4a). Early-stage activity patterns are largely uncorrelated, consistent with non-overlapping sensory pathways. However, over time, cascades converge, with activation maps becoming increasingly similar. These observations suggest that the dynamics most specific to each sensory modality occur early on and over the first several steps (see Fig. **??**). Note that these activation probability maps, which were estimated by simulating the cascade model 1000 times, represent probability that a single neuron is active at time step *t*.

**Figure 4.**
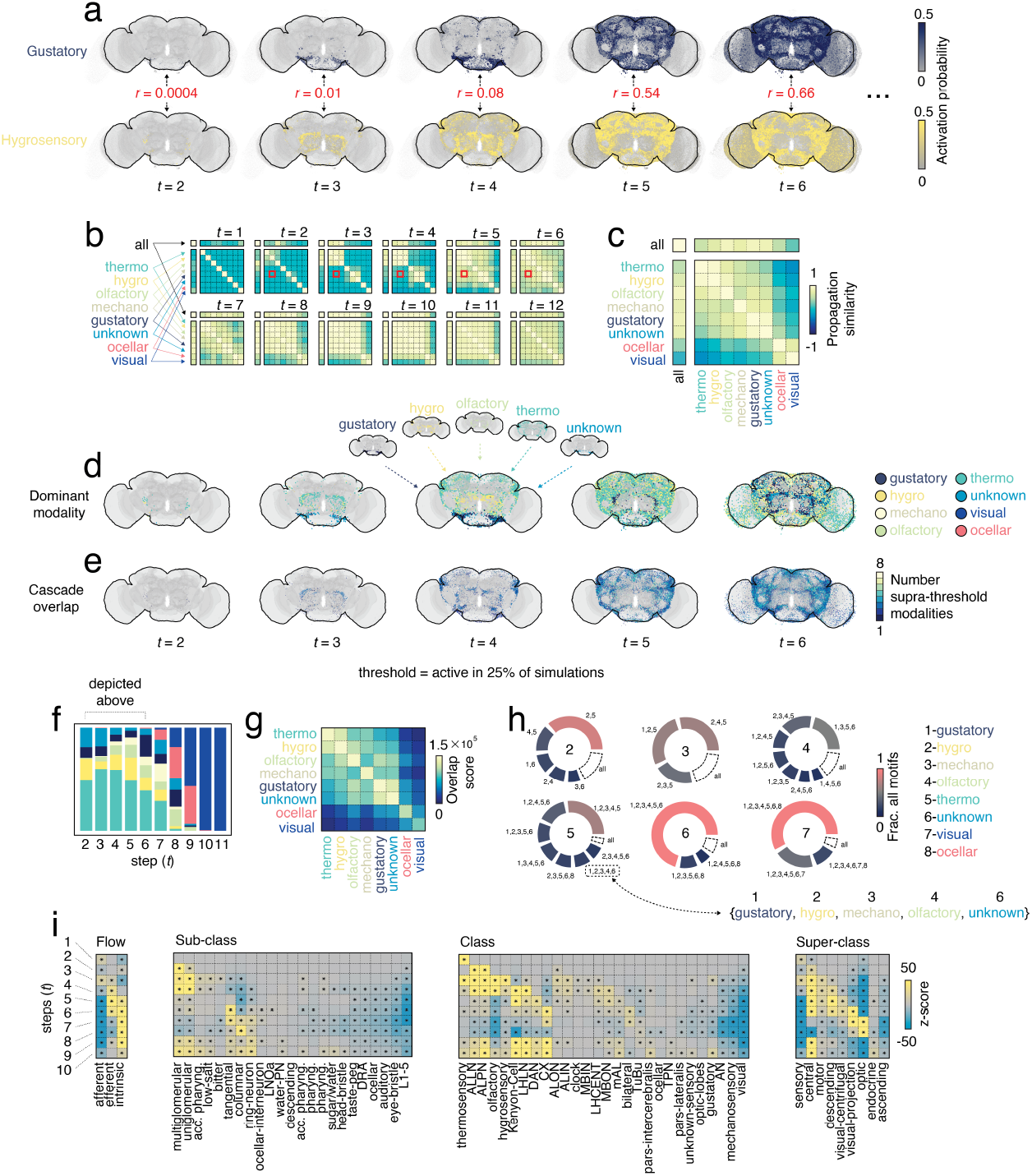
Modal specificity and similarity of cascades. Here, we consider cascades in which neurons associated with different sensory modalities are treated as seeds. From these seeds we simulate cascades and compare the spatiotemporal trajectories between sensory modalities. (*a*) Schematic for comparing cascades. For two sensory modalities (gustation and hygrosensation, in this case) we calculated the similarity between spatial maps at each time step as a correlation. (*b*) We extended this comparison to all pairs of sensory modalities as well as the “all” category (pan-sensory cascade), wherein sensory neurons from all modalities are treated as seeds, and show the similarity between all pairs at each time step, from *t* = 1 to *t* = 12. (*c*) We also concatenated the activation probability maps across time steps for a given sensory modality and computed the similarity between these omnibus maps, yielding a single time-invariant similarity score for pairs of sensory modalities. (*d*) We calculated dominance maps, where each neuron is assigned the label of the sensory modality corresponding to the greatest activation probability at a given time step. (*e*) We also calculated an “overlap index” for neurons as the number of sensory modalities with activation probability exceeding some threshold (here, 25%) at a given time step. (*f*) Composition of dominant modality as a function of time. (*g*) Pairwise overlap between pairs of sensory modalities. That is, how often did two sensory modalities simultaneously exceed the activation probability threshold together. (*h*) Frequency of overlap motifs, ranging from size 2 to size 7. (*i*) Enrichment statistics for overlap index across annotations as a function of time.

This convergence is a general feature of unimodal cascades. At early time points, each modality follows a distinct trajectory, but progressively merges into larger, more overlapping clusters (Fig. 4b). For instance, thermosensory, hygrosensory, and olfactory cascades form a cluster beginning at *t* = 2 that persists until around *t* = 5, echoing known anatomical and functional coupling between these systems [49]. Gustatory and “unknown” modalities form a smaller cluster at *t* = 3, later joined by mechanosensory inputs at *t* = 4. In contrast, ocellar and visual cascades remain distinct from all other modalities through at least *t* = 6, forming their own cluster around *t* = 7 and maintaining separation until *t* = 12.

To quantify overall relationships between modalities, we computed an “omnibus” similarity matrix. For each modality, we constructed a single vector by stacking the whole-brain activation probability vectors from time steps *t* = 1 to *t* = 12 (each vector was one-dimensional had length *N ×* 12) and calculated pairwise correlations between modalities–i.e. vectors (Fig. 4c). These relationships recapitulate known connectivity patterns—modalities with similar anatomical wiring also produce more similar cascade profiles (Fig. S3).

We further analyzed the progression of cascade overlap using two complementary statistics. First, for each neuron *i*, we identified the dominant sensory modality at each time step—i.e., the modality with the highest activation probability (Fig. 4d). Second, we calculated an “overlap index,” defined as the number of unimodal cascades exceeding a 25% activation threshold for a given neuron and time step (Fig. 4e).

Notably, dominant modalities shift over time. From *t* = 2 to *t* = 6, thermosensory cascades dominate large fractions of the network, peaking at *t* = 4 and then declining as other modalities—especially hygrosensory, olfactory, gustatory, and mechanosensory—gain influence. Visual cascades, by contrast, remain inactive until around *t* = 7, by which time other modalities are already declining.

We summarized convergence patterns by computing an overlap matrix. For each pair of modalities, we identified neurons with supra-threshold activation for both modalities at the same time step and tallied across time and space (Fig. 4g). As expected, the overlap matrix mirrors the cascade similarity matrix: visual cascades rarely co-activate with others, while thermosensory + hygrosensory and gustatory + unknown modalities frequently do. Mechanosensory and olfactory cascades exhibit less consistent clustering, suggesting more selective or delayed engagement with broader network dynamics.

To characterize multi-modal overlap more finely, we tabulated the most frequent combinations of 2 to 7 modalities co-activating a neuron. These groups consistently include thermosensory, hygrosensory, gustatory, olfactory, and mechanosensory inputs (Fig. 4h). Visual modalities appear only in the final stages, and no neurons were activated by all eight modalities under these parameters.

Finally, we performed a time-resolved enrichment analysis on overlap scores. At each time *t*, we computed mean overlap index for each annotation category, standardized against a null distribution obtained by permuting labels (Fig. 4i). As expected, early overlap of cascades is enriched among afferent, thermosensory, and sensory neurons (step *t* = 1). At *t* = 2, enrichment shifts to antennal lobe and glomerular neurons, consistent with their role in propagating activity. At later stages, overlapping cascades are enriched in intrinsic and efferent neurons (steps *t* = 3 − 5), tangential and columnar neurons (steps *t* = 6 − 7), and specific populations such as Kenyon cells, lateral horn, and central complex neurons.

Together, these results suggest that sensory cascades initially activate distinct and largely non-overlapping subnetworks, before converging on common populations of intrinsic neurons. Visual modalities remain the most distinct, engaging late and following unique trajectories through the network. These patterns reflect and reinforce known principles of modularity and sensory integration in the *Drosophila* brain.

### Cooperative sensory cascades

At the end of the previous section, we defined a measure—the overlap index—to relate the spatiotemporal trajectories of multiple sensory cascades. However, this measure was based on non-interacting cascades. A more realistic extension of the model allows cascades to interact—either cooperatively or competitively. In this sub-section, we examine a cooperative model. In this model, we seed cascades in different sensory modalities– e.g. setting *N*_*seed*_ hygrosensory neurons and *N*_*seed*_ thermosensory neurons active at time *t* = 0. Postsynaptic neurons can be activated by either modality, generally leading to earlier activation. Conceptually, this reflects polysensory integration, where multimodal sensory events (e.g., a change in temperature and visual input) occur simultaneously, boosting signal propagation speed through the nervous system. This type of early multisensory convergence has been observed *in vivo* in the *Drosophila* brain, particularly in thermosensory and olfactory circuits [49], and is consistent with integration within higher-order centers like the mushroom body and lateral horn [39, 40].

To characterize this phenomenon, we simulated cooperative cascades in which seed neurons were drawn from pairs of sensory modalities. After 1000 simulations per pair, we computed each neuron’s mean activation time and compared it to the same neuron’s activation time in unimodal simulations. We highlight these distinctions by calculating the difference between the competitive and unimodal activation probability for each sensory modality. This yielded a measure of relative speed-up—i.e., the reduction in activation time due to cooperation.

We found that, in general, cooperative stimulation resulted in modest speed-ups. Across all system pairs, the average activation time reduction was −0.11 *±* 0.05 steps. Pairs that yielded the greatest reductions included gustatory+unknown (−0.18), olfactory+unknown (−0.18), visual (retinal + ocellar) systems (−0.18), and gustatory+olfactory (−0.17) (Fig. 5a).

**Figure 5.**
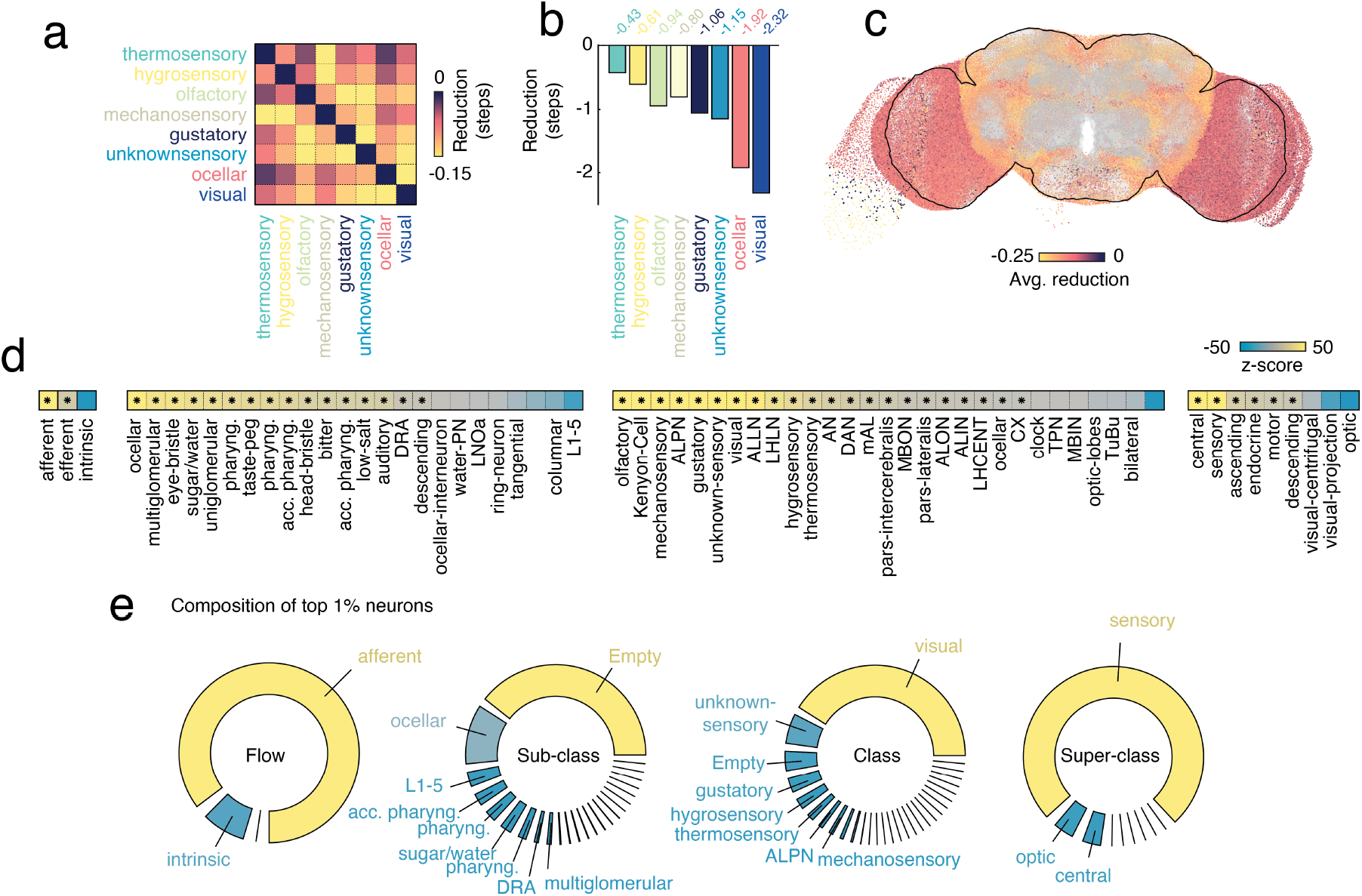
Cooperative cascade statistics. Here, we consider cascades in which neurons associated with different sensory modalities are treated as seeds. In contrast with the competitive cascade model, where postsynaptic neurons are activated by one sensory modality or the other, neurons here can be activated by *either* sensory modality. In principle, then, the cooperative cascade leads to reductions in activation time for “synergistic” sensory modalities. That is, pairs that target distinct neuronal populations or sufficiently enhance the probability of activating a single neuronal population. (*a*) Average reduction in activation time for all pairs of sensory modalities. Note that the colormap is inverted, so that the greatest reductions are depicted in bright colors. (*b*) We also compared unimodal cascade activation times against the scenario where seed neurons are selected randomly from all sensory modalities. Here, we show the mean reduction in activation time for such a scenario. (*c*) The same information at single-cell resolution and in anatomical space. (*d*) Enrichment statistics for speed-ups. (*e*) The distribution of speed-ups was heavy-tailed. Here, we highlight the tail of this distribution (the top 1% of neurons by average reduction in activation time), showing their composition across annotation categories.

To explore a more extreme case of cooperation, we simulated pan-sensory cascades where all eight sensory modalities were simultaneously seeded. This condition led to significantly faster propagation: compared to unimodal cascades originating in visual neurons, the pan-modality cascade reduced activation time by nearly 2 steps (Fig. 5b). For completeness, we visualized the reduction in activation time for each neuron in anatomical space (averaged over all eight unimodal conditions; Fig. 5c).

We also used annotation-based enrichment analyses to examine the neurons most affected by multisensory cooperation (Fig. 5d). Because activation time reductions were heavy-tailed—a small number of neurons showed large effects—we focused on the top 1% of neurons (ranked by activation time reduction). Fig. 5e summarizes the raw annotation counts for this subset, highlighting that cooperation disproportionately benefits certain neuron classes.

Together, these results suggest that visual cascades, which follow distinctive trajectories compared to other modalities, benefit most from multisensory stimulation. Nevertheless, the modest mean reductions indicate that even unimodal cascades can leverage connectome architecture to propagate efficiently through the brain.

### Competitive sensory cascades

An alternative scenario for exploring interacting cascades is the so-called competitive cascade, wherein the signal propagated from two sensory modalities retains modal specificity. That is, neurons activated by either of these two modalities inherit the “label” of their respective sensory activator, maintaining the distinct identity of each signal. For instance, we could set *N*_*seed*_ hygrosensory and *N*_*seed*_ thermosensory neurons active at time *t* = 0. The hygrosensory and thermosensory seeds are active with labels “1” and “2”, respectively. At the next time step, *t* = 1, post-synaptic neurons activated by hygrosensory neurons inherit the label “1” while those activated by thermosensory neurons inherit the label “2”. Neurons can only have one of these two labels; never both. As with the cooperative cascade, the competitive cascade model can be interpreted in terms of multisensory stimulation. However, unlike the cooperative case, where multiple sensory modalities amplify a signal, in the competitive scenario, the more “salient” of the two modalities can dominate, effectively muting responses from the other modality. This dynamic may resemble behavioral scenarios in which conflicting sensory cues must be resolved by suppressing one modality to favor another [40, 49]. In this section, we report results under this competitive scenario.

As an illustrative example, we show distinctions between unimodal and competitive cascades originating in gustatory and hygrosensory neurons. In Fig. 6a (top+bottom), we show probabilistic activation maps corresponding to unimodal gustatory and hygrosensory cascades—i.e., without any competitive interaction. However, when we allow the cascades to evolve simultaneously and compete for territory, we find distinct spatiotemporal trajectories (Fig. 6a, middle). We highlight these distinctions by calculating the difference in activation probability (competitive minus unimodal). One of the useful statistics that arises from competitive cascades is the so-called “neighborhood entropy”—a summary measure of how balanced a neuron’s neighbors are in terms of their assigned labels. A neuron whose neighbors all have only one label will have low entropy, whereas if the labels are distributed uniformly, the entropy is high. In Fig. 6b, we depict neighborhood entropy in anatomical space for the gustatory versus hygrosensory competitive cascade. Neurons with high entropy—indicated by bright colors—sit at the boundaries of competing cascades. Thus, entropy serves as a complementary metric for identifying locations where multiple signals converge onto shared targets, potentially supporting multisensory integration.

**Figure 6.**
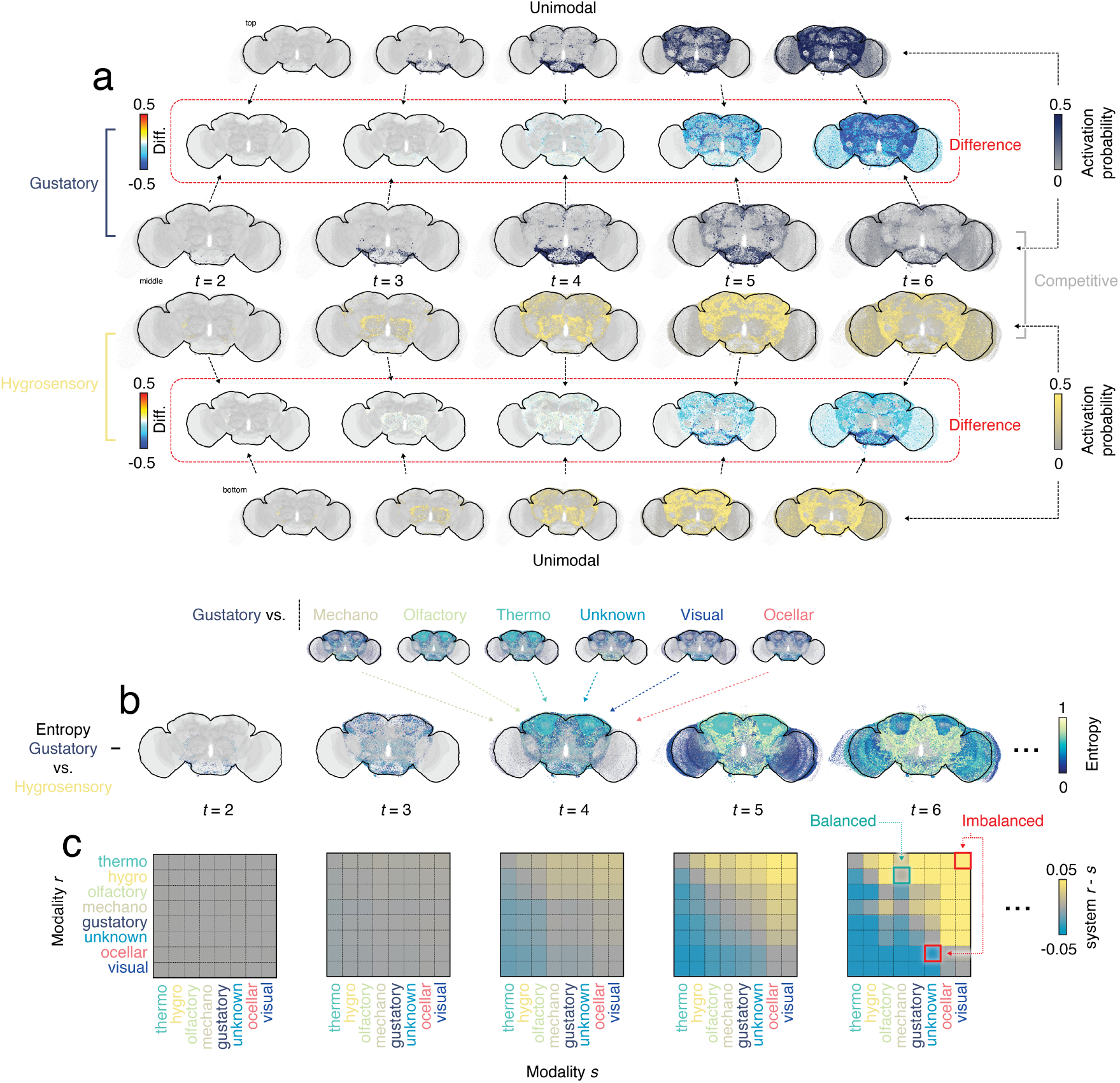
Competitive cascade statistics. (*a*) Comparison between unimodal gustatory and hygrosensory cascades (top and bottom) with competitive cascade (middle). (*b*) Entropy map for the gustatory and hygrosensory competitive cascade. High entropy neurons have “diverse” neighborhoods, i.e. their neighbors are activated by both gustatory and hygrosensory neurons. The inset with smaller brain maps depicts entropy maps of other competitive cascades involving the gustatory sensory modality at *t* = 4. (*c*) For every pair of neurons and at each time step, we calculated the relative dominance of one sensory modality over the other. That is, when we consider all active neurons, is one sensory modality over-expressed? Here, we plot this balance as a difference in proportion for all pairs of systems from time steps *t* = 2 to *t* = 6.

We estimated the “winning” sensory modality at each time step by calculating the imbalance in the number of neurons assigned to each label. Specifically, we calculate the difference in the number of neurons activated by sensory modality, *r*, compared to the other, *s* (we express this number as a fraction of all *N* neurons). In Fig. 6c, we show the balance metric as a function of time step. Each entry in this matrix corresponds to the difference in territory occupied by sensory system *r* versus *s*. For example, at *t* = 6, two cascade pairs show strong imbalances (red). In one case, thermosensory neurons outcompete visual neurons; in another, unknown sensory neurons dominate ocellar inputs. We also show a more balanced case (hygrosensory *vs*. mechanosensory), where territorial differences are minimal at *t* = 6.

This competitive cascade model helps us better understand how multisensory signals might resolve within a weighted and directed connectome. For instance, we can use neighborhood entropy to track the formation of “fronts” or boundaries between competing sensory signals. Neurons with high entropy are structurally poised for integration—they receive input from, or project to, neurons with differing sensory identities. Because the *Drosophila* connectome is directed, we defined three forms of entropy: based on presynaptic partners, postsynaptic partners, and their union. In a pan-sensory competitive cascade model, where all sensory systems were seeded equally, these three measures were highly correlated (mean pairwise similarity across time points *r* = 0.81 ± 0.21). Nonetheless, subtle differences were evident, likely due to the strong—but not perfect—correlation between in- and out-degree (*r* = 0.82) [39].

We further found that a neuron’s degree was strongly predictive of its time-averaged entropy (*r* = 0.91; Fig. 7b). In general, high entropy neurons tended to be structural hubs, consistent with their broad connections across sensory domains. This was true for both binary and weighted degrees (Fig. 7c). However, a neuron’s role in sensory integration is not static: entropy varied across time. Mean entropy increased rapidly to a peak at step 5 before decreasing (Fig. 7d). Entropy maps showed low similarity across time—maps from early and late steps were even anti-correlated (Fig. 7e), suggesting a complex temporal dynamic shaped by cascade propagation and network topology.

**Figure 7.**
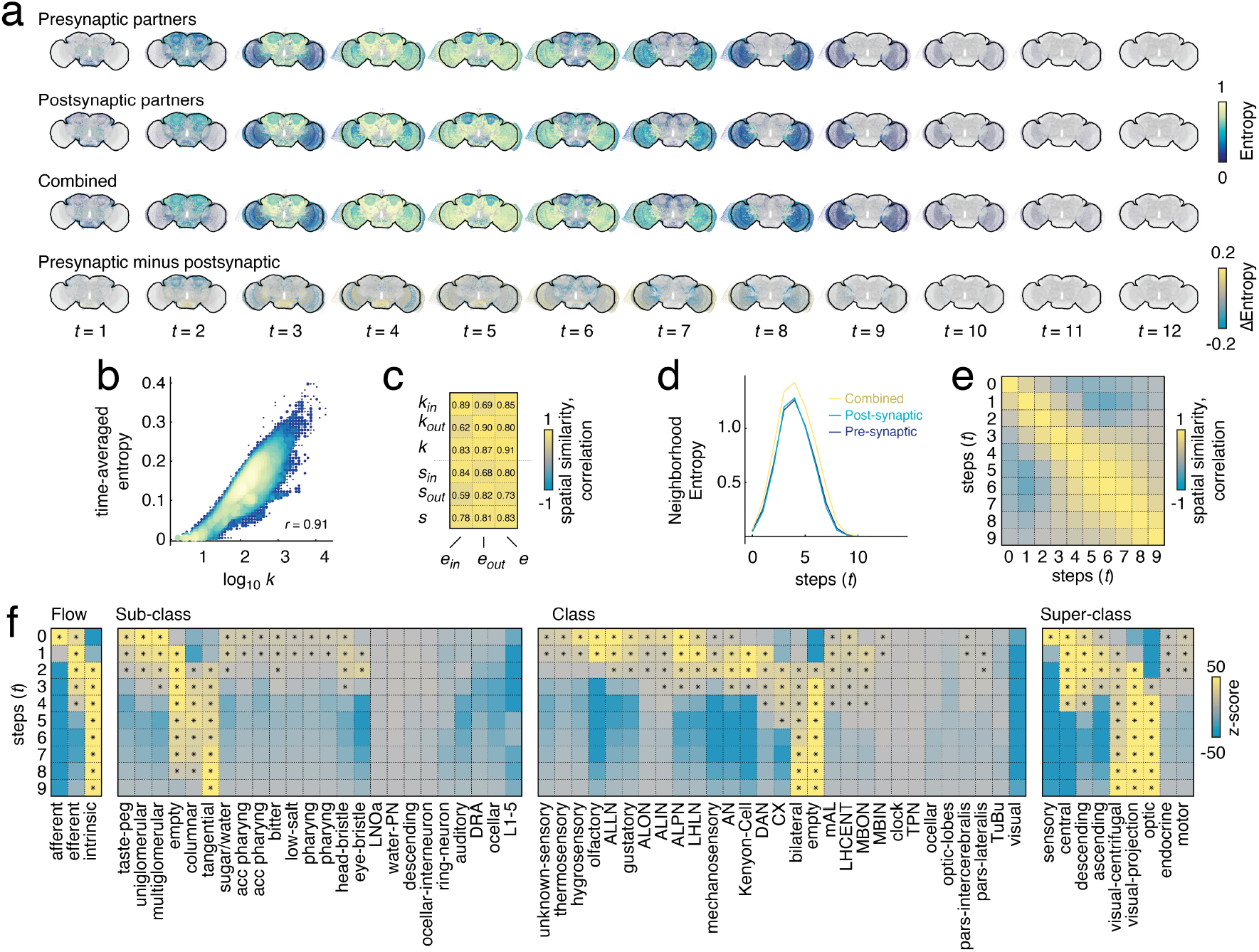
Additional competitive cascade statistics. The entropy patterns vary depending upon how one defines a neuron’s “neighborhood”. For instance, one can define a neighborhood based on a neuron’s pre- or post-synaptic neighbors or the combined neighborhood. Here, we compare these three definitions in panel *a*. We also related entropy to conventional network statistics. (*b*) Scatterplot of time-averaged entropy against the logarithm of nodes’ total degrees. (*c*) Correlation of entropy with various other network statistics: in/out/total degree and strength. (*d*) Mean whole-brain entropy of different neighborhood definitions as a function of step. (*e*) Similarity of whole-brain combined-neighborhood entropy patterns between time steps *t* = 0 to *t* = 9. (*f*) Enrichment statistics for whole-brain combined-neighborhood entropy.

Finally, we examined whether neighborhood entropy was enriched within specific cell types and brain regions. As in previous analyses of activation time and overlapping cascades, we found distinct temporal trajectories by annotation. Afferent sensory neurons peaked in entropy early (steps 1–2), followed by efferent and intrinsic neuron types—including columnar neurons, Kenyon cells, central complex neurons, ascending/descending neurons, and visual centrifugal neurons (Fig. 7f). These trajectories reinforce earlier observations that central brain neurons serve as hubs for signal convergence [39, 40], with their entropy dynamics reflecting their role in integrating competing signals over time.

These observations suggest that intrinsic neurons in the central brain are key loci of sensory integration. Positioned at the boundaries of competing cascades and exhibiting high neighborhood entropy, these neurons are structurally and functionally poised to arbitrate between conflicting or convergent sensory inputs. Although high-degree neurons tend to show high entropy, the temporal dynamics of the competitive model underscore that a neuron’s role as integrator depends on timing and network context, not just connectivity.

### Data-driven clustering of neurons based on cascade properties

We previously introduced a set of summary statistics derived from the spatiotemporal evolution of sensory cascades. These included activation time, overlap, competitiveness (entropy), and cooperative speed-up. We used enrichment analyses to relate these features to neurobiological annotations. Here, instead of aligning with known labels, we clustered neurons based solely on cascade-derived features to identify data-driven functional groupings.

To do so, we aggregated the aforementioned statistics—overlap, entropy (from competitive cascades), reduction in activation time (from cooperative cascades), and activation timing—into an *N* × *N*_features_ matrix. We standardized (z-scored) each feature and applied *k*-means clustering using a correlation-based distance metric (Fig. 8a). We varied *k* from 1 to 50, performed 100 random restarts per *k*, and selected *k* = 6 based on the “elbow” of the within-cluster sum of squared errors curve (Fig. 8b). Spatial projections of these six clusters are shown in Fig. 8b.

**Figure 8.**
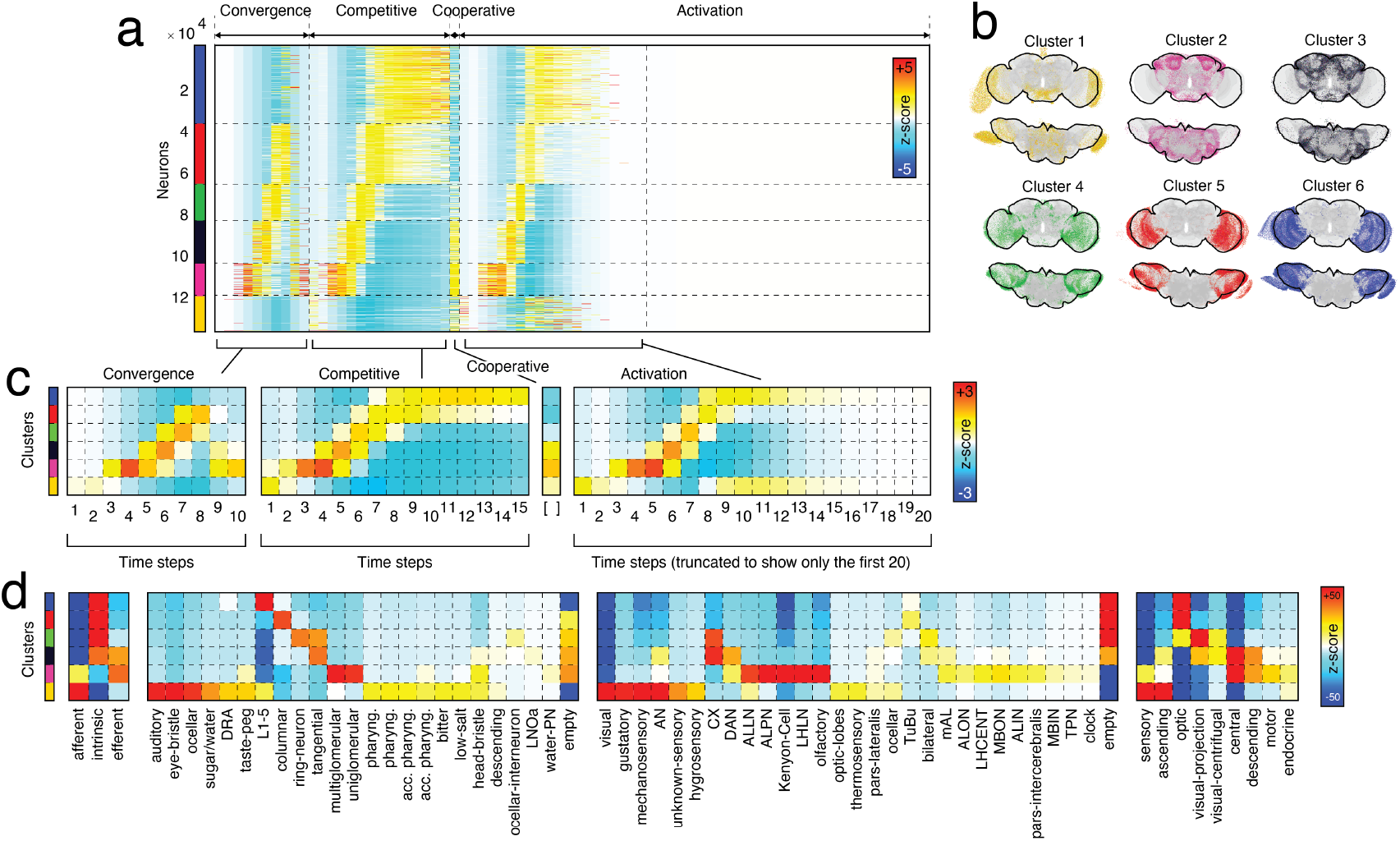
Clustering neurons based on their cascade properties. (*a*) Neurons ordered by cluster assignment. Columns correspond to z-scored cascade metrics. (*b*) Cluster labels depicted in anatomical space. (*c*) Cluster centroids. (*d*) Enrichment of cluster labels across annotations. Note that at the far right of panel *a*, we show the activation probability for each neuron at time points 1-50. In panel *c*, we show cluster centroid time courses for steps 1-20. Note further that for the purposes of clustering, we do not use the full activation time courses. Rather, we summarize the neuron *×* time point matrix into a neuron *×* 1 vector whose entries denote the average time of activation for each neuron.

Each cluster was associated with a distinct centroid—an archetypal profile across cascade features (Fig. 8c). Cluster 1 (yellow), for example, exhibited the earliest activation times and early-onset entropy. This cluster was enriched for afferent, sensory, and ascending neurons, consistent with its role as a cascade initiator. Cluster 2 (pink) followed temporally, with peaks in overlap, entropy, and activation between steps *t* = 3 and *t* = 6. It was enriched for efferent neurons, multiglomerular/monoglomerular projection neurons, and central complex interneurons, including Kenyon cells.

Clusters 3 through 6 exhibited progressively later activation profiles and distinct enrichment patterns. All were composed largely of intrinsic neurons and converged on visual projection and centrifugal neurons in the optic lobes.

These findings suggest that sensory spreading dynamics constrained by the connectome can be used to derive functionally meaningful, temporally ordered groupings of neurons. These groups reflect similarity in cascade participation and their respective roles in competition, cooperation, and convergence across the network.

### Additional analyses

We conducted several supplementary analyses to assess the robustness and generality of our results. First, we verified that our findings were not sensitive to weak synaptic connections: thresholding the connectome to retain only synapses with at least five connections had minimal effect on results. Summary statistics remained highly correlated with those from the unthresholded network (Spearman correlations: *ρ*_act_ = 0.80, *ρ*_coop_ = 0.56, *ρ*_comp_ = 0.82; all estimated with *p*_transmission_ = 0.01 and *N*_seed_ = 16).

Second, we showed that activation timing could not be predicted solely from shortest path length—a static graph measure derived from connectivity alone (Fig. S5; see also [50]). Third, we examined how cascades unfolded across neurotransmitter classes, revealing transmitter-specific activation time distributions (Fig. S6).

Fourth, we compared sensory-driven cascades to cascades seeded from random non-sensory neurons. These exhibited systematically different spatiotemporal profiles, underscoring the distinctive topological embedding of sensory neurons (Fig. S7).

Fifth, we analyzed community structure in the connectome using a nested stochastic block model [51, 52]. Cascade-derived statistics were highly enriched within and segregated between communities (Fig. S8, Fig. S9). These findings support the idea that communities serve as mesoscale substrates for sensory integration and signal flow (see also [53–55]).

As a complementary continuous approach, we performed a principal components analysis on cascade features. The resulting components captured brain-wide gradients of variation and reflected global organizational motifs in cascade dynamics (Fig. S10).

## DISCUSSION

The *Drosophila* connectome contains approximately 140,000 neurons and over 15 million synaptic connections, forming a sparse yet structured network with profound implications for information flow and processing. As we showed, sensory neurons from different modalities are highly segregated, with minimal direct synaptic links, suggesting architecture tailored for specialized, parallel sensory processing.

Yet many brain functions require integration across signals—such as decision-making [56], navigation, or motor planning—and depend on multi-step polysynaptic pathways not obvious from first-order connectivity. One way to explore these circuits is through dynamic simulations constrained by the connectome, rather than relying solely on anatomical graphs [57, 58].

In this work, we employed a spreading, cascade-style model that abstracts away full biophysical detail in favor of analytic clarity. While mechanistic models (e.g., integrate-and-fire) offer realism, our approach foregrounds interpretability, enabling us to trace cascade evolution directly under network constraints. Related spreading models have informed human brain dynamics [27], as well as larval *Drosophila* modeling [43]. Our results show that sensory cascades—initially modality-specific—rapidly converge onto overlapping sets of secondary nodes, many located in the central complex and involving descending and motor neurons. This proximity between sensors and effectors is consistent with behavioral observations and suggests streamlined sensorimotor pathways.

Of course, the simplicity of our model introduces limitations. Cascades always reach an inactive equilibrium, which is biologically unrealistic, yet this property enables clear ranking of neurons by mean activation time—a metric tightly controlled by user-defined seed size and transmission probability, and directly tied to connectivity. Despite its outward simplicity, the application of our model to a directed and weighted, neuron-resolved connectome represents an important conceptual advance. Previous applications to human connectome data focused on undirected connectomes [27] and applications to directed networks have been limited to meso-scale, interareal connectomes [29]. A natural target for future studies concerns understanding precisely how the connectome’s scale (neuron resolution versus interareal) and directedness shape the spread of activity.

Comparatively, computational studies using whole-brain leaky integrate-and-fire models (e.g. [59]) demonstrate excellent alignment with experimental data for gustatory and grooming circuits. Those bottom-up models focus on detailed circuit dynamics and specific behaviors, while our top-down network approach accommodates broad hypotheses about sensorimotor flow from minimal assumptions.

We offer testable predictions around multisensory integration—specifically that thermo- and hygrosensory pathways co-activate overlapping neuronal groups, as identified by our overlap index. These hypotheses can be evaluated experimentally using paired or combinatorial multisensory stimulation paradigms. For instance, using the cascade model, we could identify multi-sensory hub neurons that are consistently coactivated by two different modalities. If sensory information from those modalities was relevant for a particular behavior, we might hypothesize that perturbing the predicted hubs would impact that behavior more negatively than, say, a population of spatially co-localized non-hub neurons.

Another important insight comes from the passive nature of our model: activation flows without external control signals, emerging solely from structural connectivity. This emphasizes the role of network architecture in shaping dynamic propagation—even without directed regulation.

Looking forward, combining structure with experimental data—e.g. matching stimulus-evoked recordings to connectomic projection maps—offers promising opportunities for linking wiring to function more directly. Similar approaches in human macroscale network neuroscience have already yielded predictive insight about functional connectivity [60, 61].

Our model could be further enhanced by including synaptic polarity, inhibitory/excitatory distinctions, and transmitter-specific dynamics. For instance, connectome-informed weight estimates often disagree with simply assuming synapse count corresponds to weight [62]. Likewise, expanding the model to include ventral nerve cord circuits and motor neuron pathways would better approximate the full sensorimotor axis.

Another strategy for making the cascade model richer and more biologically realistic is to allow for asynchronous updating. That is, rather than forcing all neurons to transition synchronously at the same step, allow signals to propagate through space, such that a presynaptic neuron can influence its post-synaptic partner faster if those two neurons are closer to one another. Although appealing, this type of asynchronous updating presents a number of challenges. First, the synchronous model studied here, despite its outward simplicity, is quite complex; added modeling components, e.g. asynchronous updating, may create lead to interpretational challenges at the expense of relatively little gain in biological realism. Second, implementing the asynchronous updating would be difficult due to the polyadic connectome. That is, a pre- and post-synaptic neuron can connect *via* multiple synapses with distinct locations along their respective processes. From a modeling perspective, this means that the delay associated with a signal traveling to the soma of the post-synaptic neuron is variable depending on the specific synapse through which it was transmitted. This complication is difficult to implement largely due to the scale of the synapse-resolved connectome; previous studies have incorporated delays using meso-scale connectome data [27, 29] in which the connectomes were orders of magnitude smaller and for which pairs of brain areas were connected by only a single connection. We leave this extension for future studies.

There are also other (less challenging) modeling issues that could be addressed in future studies. Here, despite vast differences in the number of neurons associated with each sensory modality (e.g. roughly 10^4^ visual neurons but 10^1^ thermosensory neurons), we forced the number of seeds to be constant across modalities, ensuring balance. On the other hand, keeping the number of seed neurons constant impacts the consistency of cascades from one simulation to the next. For instance, with *N*_*seed*_ = 16, there is always overlap in the set of seeded neurons for thermosensory cascades; for the visual modality, it is unlikely that any two cascades are initialized with the same set of seeds. This likely has a noticeable effect on some of the measures we consider– e.g. cascade overlap and dominance maps, which require a neuron to be active in 25% of simulations to be considered. If the future, one could consider informing this parameter on the basis of experimental evidence– e.g. when presented with a visual stimulus, roughly what fraction of retinal neurons are active?

The cascade models studied here also connect, albeit imprecisely, with other phenomena observed in neural systems, including wave patterns. Wave-like phenomena in brain imaging data–e.g. traveling, standing, and colliding waves [63]–can be viewed as forms of activity spreading across the connectome [64]. Much like cascades in network science [65], these waves propagate sequentially along structural connections, depend on local neighborhood interactions, and show amplification and attenuation. Both are strongly shaped by the same underlying connectome: hubs, gradients, recurrent loops, and the connectome all influence how activity flows. Relevant to our work here, interactions between multiple waves can even resemble competing cascades.

However, neural waves are richer than typical cascades. They are continuous spatiotemporal fields governed by neural biophysics—including conduction delays, excitation–inhibition balance, and oscillatory stability—where geometry matters and interference or superposition can occur. Cascade models, like the ones we study here, usually treat nodes as discrete on/off units and lack superposition or oscillatory behavior. Still, a unifying perspective is that neural waves can be understood as cascade-like spreading processes operating on weighted, continuous, and delay-coupled graphs, making the two formalisms closely related but not identical. A critical direction for future work involves directly connecting activation statistics to decision variables, establishing a clearer correspondence between cascades and behavior [66]. For instance, we could treat premotor, descending neurons as read-out neurons that project to specific body segments or effectors (wings, limbs), enabling specific behaviors [67, 68]. Doing so may also allow us to generate more meaningful summary statistics, using relevant aspects of dynamics to contextualize the importance of architectural features of the connectome [69].

One caveat remains: the current *Drosophila* connectome is incomplete—roughly half of all synapses are unassigned [35]. That said, our sensitivity analysis using a thresholded connectome (only edges with ≥5 synapses) produced similar activation ordering, with generally slower timescales. This suggests increased density from filling missing synapses would mostly compress timescales, preserving relative ranking among neurons.

More broadly, our work bridges nanoscale connectomics with system-level network neuroscience. While most network theory has focused on meso- and macro-scale graphs revealing small-world structure, rich clubs, modularity, and cost-efficiency, we demonstrate that the same tools and principles apply—and even reveal new patterns—at the synapse-level resolution. Unlike previous work in larger brains, our cascades uncover functional trajectories and temporal motifs that arise purely from synaptic architecture.

Taken together, this suggests that synapse-scale connectomes may shape—and be shaped by—the same organizational principles operating in larger neural networks, while also supporting their own novel dynamic motifs specific to sensorimotor processing in small, tractable nervous systems.

## MATERIALS AND METHODS

### Drosophila connectome dataset

All connectome data, including synaptic connectivity, three-dimensional coordinates, neuropil volumes, and annotations were obtained from https://codex.flywire.ai/. These derivatives represent the output of a process beginning with nanometer-resolution electron microscopy images of an adult female *Drosophila* [70]. The FlyWire project allowed citizen scientists to trace neurons [71] to reconstruct the connectome. Combined with automated approaches [72, 73], the end result was a complete map of synaptic connections among approximately 139,000 neurons.

Neurons in *Drosophila* are polyadic, thus the same pair of neurons can synapse onto one another at multiple distinct sites. The https://codex.flywire.ai/ are shared in two versions: one in which pairs of neurons were connected if they were linked by, at minimum, five synapses and another in which even a single synapse was considered evidence that two neurons were connected (unthresholded). In the main text, we analyze the unthresholded connectome, but note that results are consistent with those obtained using the thresholded version.

Annotation data were obtained from at the same time as the connectome data. Annotations were grouped into four distinct categories [37]. The first two annotations are “dense,” in the sense that every neuron is assigned a label.

1. “Flow”: afferent, efferent, intrinsic.
2. “Super-class”: sensory (periphery to brain), motor (brain to periphery), endocrine (brain to corpora allata/cardiaca), ascending (ventral nerve cord (VNC) to brain), descending (brain to VNC), visual projection (optic lobes to central brain), visual centrifugal (central brain to optic lobes), or intrinsic to the optic lobes or the central brain

The other two categories are not dense–i.e. not all neurons are assigned a label. The “Class” and “Sub-class” annotations are nested hierarchically within the the parent “Super-class” and describe cell groups reported in the extant literature, e.g. neurons in the optic lobe [74].

### Cascade model

Here we explore an abstract cascade model. In this model, each neuron exists in one of three states: active, inactive, or refractory. In our simulations, we set select sets of neurons to be active at time *t* = 0. These initially activated neurons are referred to as “seeds” and are selected from among sensory neurons alone. The seeds are intended to represent sensory stimuli.

In this model, neurons that are active at time *t* probabilistically activate their post-synaptic partners at *t*+ 1. Suppose neuron *i* makes a single synapse onto neuron *j*. Given that *i* is active at *t*, the probability that neuron *i* activates *j* at *t* + 1 is given by the transmission probability parameter, *p*_*transmission*_. Conversely, the probability that *i* does *not* activate *j* is simply *q* = 1 − *p*_*transmission*_.

Oftentimes, neuron *i* makes multiple synapses onto *j*. We treat these as independent trials; only one successful trial is sufficient to activate neuron *j*. What is the probability that at least one event is successful? In the space of possible outcomes, there is exactly one outcome in which none of the synapses successfully activate *j*. Its probability is given by: *q*^#*synapses*^ = (1 − *p*_*transmission*_)^#*synapses*^. All other possibly outcomes include at least one successful transmission event. Hence, the probability that *j* is activated by neuron *i* at time *t* + 1 is given by: 1 − *q*^#*synapses*^.

For all unimodal, cooperative, and competitive cascades, 16 neurons from each sensory modality were selected randomly as seeds for the cascades.

At a high level, we view each cascade a single “sensory event” passing through the network. Previous studies have modeled this type of event using only “inactive” and “active” states (as in previous studies, e.g. [27, 29]). That is, once a neuron is active, it stays active for the duration of the cascade and can activate its post-synaptic partners at every step after it becomes active. Here, we opt to introduce a third “refractory” state. Neurons enter this state immediately the step immediately following their activation.

The sequence “inactive→active→refractory” is consistent the simplest physiological spike cycle. The refractory state, which distinguishes our work from other related studies on cascade models, is appropriate for the timescales of sensory events, whereby channel inactivations following a spike prevent neurons from spiking again until recovery (which we assume takes much longer than the timescale of a sensory event). Moreover, this state sequence helps to create unidirectional waves of activation, which are consistent with observations of propagating calcium waves in Drosophila [75, 76] and zebrafish [77].

Synaptic activation in our cascade model is implemented as a Bernoulli process in which each structural synaptic contact provides an independent opportunity for postsynaptic activation. This formulation has a direct biological interpretation: real chemical synapses exhibit low and probabilistic vesicle-release rates, with measured release probabilities commonly ranging from 1–10% at many central synapses [78, 79]. Moreover, individual synaptic contacts often behave as independent release sites, consistent with classical quantal models [80] and with ultrastructural reconstructions in Drosophila showing multiple discrete active zones per connection [81, 82]. The effective probability that a presynaptic spike triggers a postsynaptic spike within a brief cascade timestep is further reduced by dendritic integration constraints [83], inhibitory shunting [84], short-term synaptic depression [85], and spike-threshold variability [86]. Thus, small transmission probabilities (1–5%) are biologically realistic when interpreted as the net probability that a single structural synapse produces a downstream spike during a transient propagation event.

In our model, cascade dynamics are initiated by activating a small number (*N*_*seed*_) of seed neurons at *t* = 0, which reflects the fact that many neural events begin with sparse, localized activation rather than widespread population firing. Sensory pathways frequently exhibit highly selective and sparse initial responses, as seen in Drosophila olfaction where only a small subset of receptor neurons and projection neurons respond strongly to a given odor [87, 88] and in vertebrate cortex where sparse coding is a dominant organizational principle [89]. Sparse initial activation is also characteristic of feature-selective neurons in early sensory systems [90] and of startle and escape circuits, where a very small number of command-like neurons seed fast recruitment [91]. Moreover, large-scale neural events such as cortical traveling waves or hippocampal sharp-wave ripple events typically originate from small focal clusters of neurons before propagating across the network [92, 93]. Thus, seeding the cascade with a small set of neurons captures the biological reality that many propagating population events begin from a minimal, spatially or functionally localized trigger, consistent with both sensory-evoked activity and spontaneously initiated neural dynamics.

#### Activation probability

The cascade model is stochastic. In principle, each run can yield a different sequence of neuronal activations. For every experiment, we summarize across 1000 independent runs by estimating the probability a given neuron was active at time *t* (see Fig. S1 for example maps). We denote the activation probability of neuron *i* at time *t* as *P*_*i*_(*t*). In the main text we describe several statistics calculated directly from the probabilistic activation curves.

#### Dominance maps

We use the cascade model to simulate sensory cascades by allowing a fixed number of sensory neurons with specific modalities to be active at time *t* = 0. In total, we consider eight sensory modalities as determined by hierarchical annotations of sensory neurons: thermosensation, hygrosensation, olfaction, mechanosensation, gustation, vision (ocellar and compound), and an unknown sensory class (no clear assignment). For estimate modality-specific activation probability curves for each neuron. Given that sensory neurons associated with modality *m* were seeded at time *t* = 0, we denote the probability that neuron *i* is active at time *t* as: 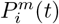. From these estimates, can calculate the “dominant” modality for each neuron at any given timepoint as the *m* that maximizes 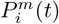.

#### Overlap score

The dominance index represents a “winner take all” measure, ascribing dominance to a single sensory modality. In reality, there could be multiple near-maximal modalities, all with similar activation probabilities. To account for this possibility, we defined the overlap index. Specifically, we define a threshold, *τ*, and calculate: 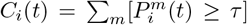. That is, we count the number of sensory modalities for which the activation probability of node *i* at time *t* is at least *τ*.

#### Cooperative cascades

We explored several variations of the cascade model. The first variant focused on “unimodal” cascades, wherein the seed neurons were selected from among a single sensory modality. However, we also considered a second “cooperative” variant, in which pairs of sensory systems were seeded (doubling the total number of seed neurons). We then calculated the reduction in the mean activation time (speed-up) as a result of the dual-seed approach as: *A*_*rs*_ − min(*A*_*r*_, *A*_*s*_), where *A*_*rs*_ is the activation time of a given neuron under the cooperative, dual-seed model *A*_*r*_ and *A*_*s*_ are the activation times of the same neuron under the unimodal model where modalities *r* and *s* were seeded independently.

#### Competitive cascades

We also considered a “competitive” variant of the cascade model. In this case, we imagine two distinct sensory signals (“blue” and “red”) spreading throughout the connectome carrying distinct information. Postsynaptic neurons activated by a majority blue/red presynaptic neurons will be blue/red at the following timestep. In the rare event of a tie–i.e. equal number of red and blue activating neurons–the post-synaptic neuron becomes blue/red at the following timestep with equal probability.

#### Neighborhood entropy

Under the competitive model, we can calculate an entropy measure that describes whether a node is positioned in the network so that its neighbors are mostly homogeneous in terms of their redness/blueness or whether their neighbors are diverse. To do this, we identify each neuron’s connected neighbors and, for at time *t*, those the proportion of those that are active and are blue/red. We then rescale these values so that their total sums to 1, and calculate an entropy, *H*_*i*_(*t*) = −*p*_*blue*_*log*_2_(*p*_*blue*_)+ −*p*_*red*_*log*_2_(*p*_*red*_). Intuitively, when this entropy is close to 1, it implies that the distribution of blue/red neighbors is approximately equal. However, values close to zero imply that one of the two colors is more dominant.

An important consideration is how one defines “neighborhood” in this context. In the main text, we consider three possibilities: a neuron’s pre-synaptic partners, its post-synaptic partners, and the union of those two sets.

Note that in all of the above calculations, we disregard inactive or quiescent neurons. That is, only active neurons contribute to the entropy scores.

### Community detection

Many studies have shown that biological neural networks at varying spatial scales can be meaningfully partitioned into clusters or communities [5, 52, 94–101]. In general, community structure is unknown ahead of time and must be estimated or inferred algorithmically. The procedure is referred to as community detection [102], and there exist many approaches for doing so [103–108], each of which makes specific assumptions about what constitutes a community.

One of the oldest approaches for community detection is the stochastic blockmodel [109–113]. Briefly, blockmodels imagine that every node in the network is assigned a label. The probability that any pair of nodes are connected and the weight of that connection depends only upon the communities to which they are assigned. In general, blockmodels estimate community labels and connection probability/weight distributions so as to maximize the likelihood that the model generated the observed network.

Recently, there has been renewed interest in the application of blockmodels to brain network data [20, 100, 114–121]. Here, we leverage a nested and weighted variant of the classical blockmodel to partition the *Drosophila* connectome into hierarchical communities [51]. Using this model, we obtained an estimate of the consensus hierarchical partition using a Markov Chain Monte Carlo procedure [122]. Briefly, this procedure entails allowing the Markov chain to equilibriate and sampling 10000 high-quality partitions. These partitions are aligned to one another to discover latent “modes” – partitions that share many characteristics with one another [123]. From these modes, we obtained for each neuron the community it was most likely assigned to. All analyses using communities were carried out on the resulting partition (Fig. S8 and Fig. S9).

### Clustering based on cascade properties

In the main text, we derived a series of statistics for neurons based on their activation times, overlap, neighborhood entropies, and speed-ups. We can use these data to partition neurons into clusters based not on their connectivity (information derived from the connectome) but these dynamic, cascade-based properties.

To do this, we constructed a [*N × F*] matrix, where *N* is the number of nodes and *F* = 27 is the total number of features. In this case, we considered the following features derived from omnibus sensory cascades–i.e. where all sensory modalities were seeded:

1. **Activation probability**: The [*N ×* 1] vector of each neuron’s mean activation time.
2. **Overlap index**: The [*N ×* 10] array of neurons’ overlap indices over the first 10 time steps.
3. **Neighborhood entropy**: The [*N ×* 15] array of neurons’ neighborhood entropy (estimated using both pre-/post-synaptic partners).
4. **Speed up**: The [*N ×* 1] vector of speed-ups (reductions in activation times) from single-modality unimodal cascades to the omnibus cascade.

To cluster these data, every column was standardized (z-scored). We then used k-means clustering to assign neurons to a unique cluster [124]. We selected the optimal number of clusters based on the inflection point in the summed square errors of cluster centroids relative to their constituents. Based on these analyses, we identified *k* = 6 as the optimal number of clusters.

### Annotation enrichment analysis

Throughout this manuscript we describe an “enrichment” procedure for linking continuous data to categorical labels–e.g. annotations. The aim of this procedure, in more detail, was to determine if a particular variable was over-expressed (enriched) within a given annotation category. To do this, we calculated the variable of interest’s mean value within all neurons associated with a specific annotation. We then randomly permuted the annotation labels (keeping the total number constant) and calculated the new mean. We repeated this procedure 1000 times, generated a null distribution of mean values. The enrichment score for that variable within that annotation category is the original mean expressed as a z-score with respect to the null distribution.

## Data and code availability statement

All data are publicly available through https://codex.flywire.ai/ with no restrictions. Code is available upon request.

---

**Figure S1.**
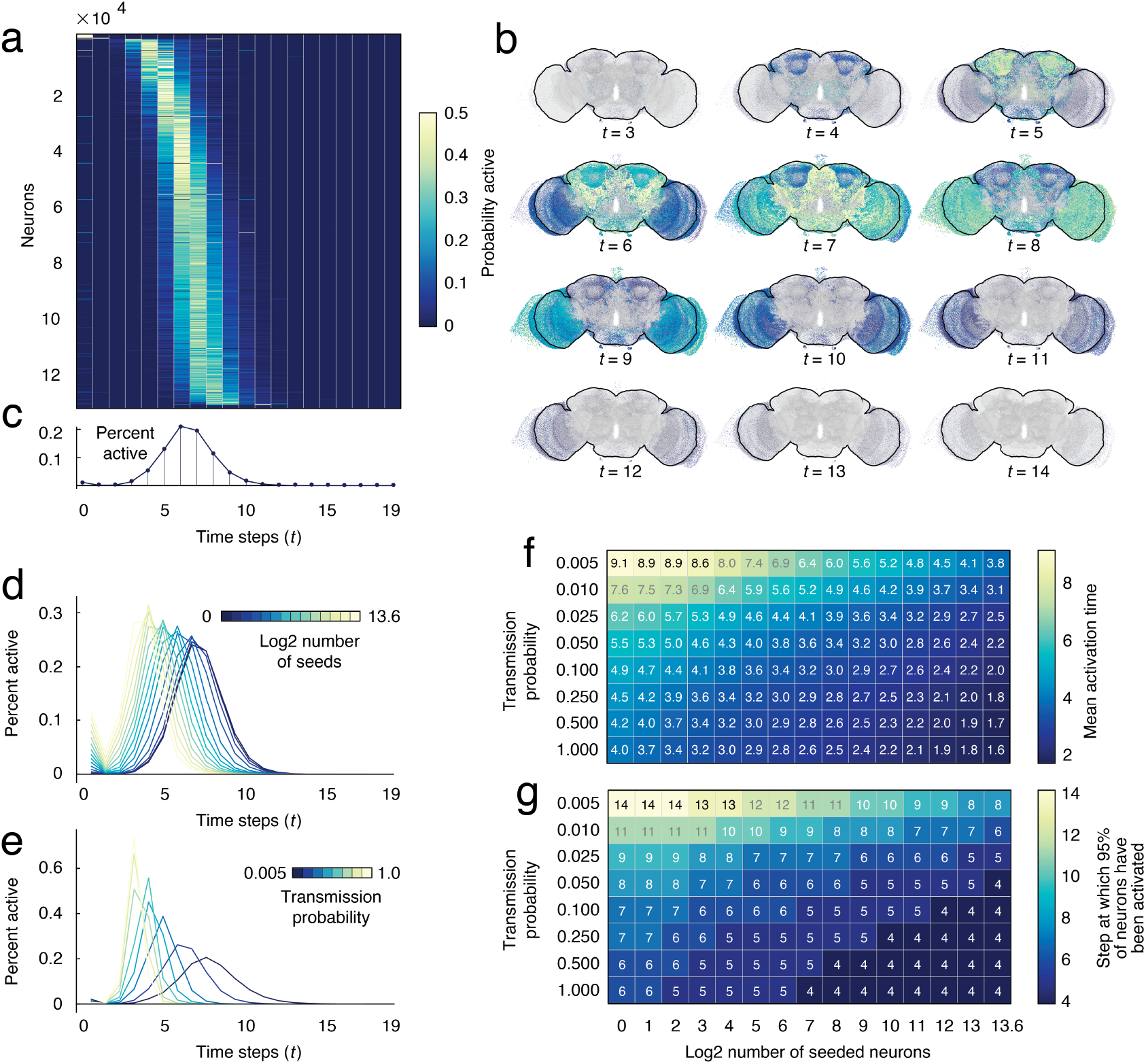
Unimodal cascades. We initialize cascades by “seeding” sensory systems with a subset of active neurons. These neurons probabilistically activate their post-synaptic neighbors. (*a*) Activation probability for all neurons × time. In this example all sensory neurons were active at time *t* = 0. (*b*) Percent of neurons active at each time step. (*c*) Activation probability of each neuron in anatomical space. Note that here we only plot each neuron’s soma and omit their arborization. The cascade model depends on two parameters: the number of seed neurons (those active at time *t* = 0) and the probability of transmission *via* a single synapse. Panels *d* and *e* illustrate how variation of these parameters impact transmission probability curves. (*d*) Increasing the number of seed neurons largely shifts the curve to the left. In this example transmission probability was held constant at *p*_*transmission*_ = 0.01. (*e*) Increasing transmission probability compresses and shifts and the curve. Panels *f* and *g* show the mean activation time for cascades at different parameter values and the mean step at which 95% of all neurons were activated, respectively. The latter measure serves as a rough estimate of when cascades end.

**Figure S2.**
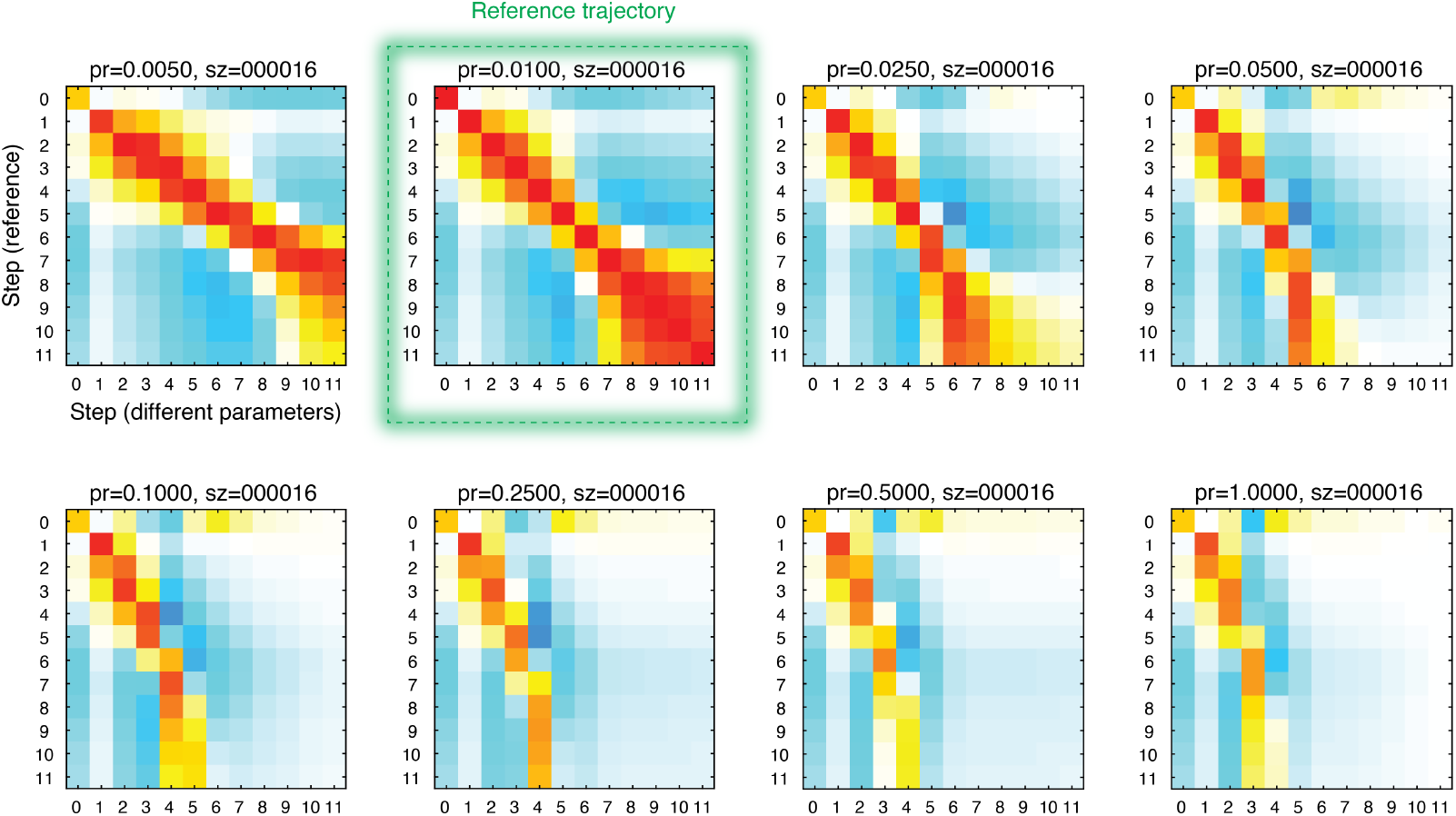
Similarity of probabilistic cascades across values of *p*_*transmission*_. In the main text, we reported results with cascade parameters fixed to values of *p*_*transmission*_ = 0.01 and *N*_*seed*_ = 16. At each time step, this “reference trajectory” (shown outlined in green in the above figure) yields a *N* × 1 vector at each time step whose *i*th entry corresponds to the probability that neuron *i* was active at that instant. To demonstrate that varying the parameter *p*_*transmission*_ generally compresses or stretches the trajectory (while only subtly changing the activation patterns themselves) we compared the reference trajectory against trajectories generated when *p*_*transmission*_ = [0.005, 0.025, 0.05, 0.1, 0.25, 0.5, 1.0]. In general, we found high degrees of spatial similarity and sequencing across these parameter values.

**Figure S3.**
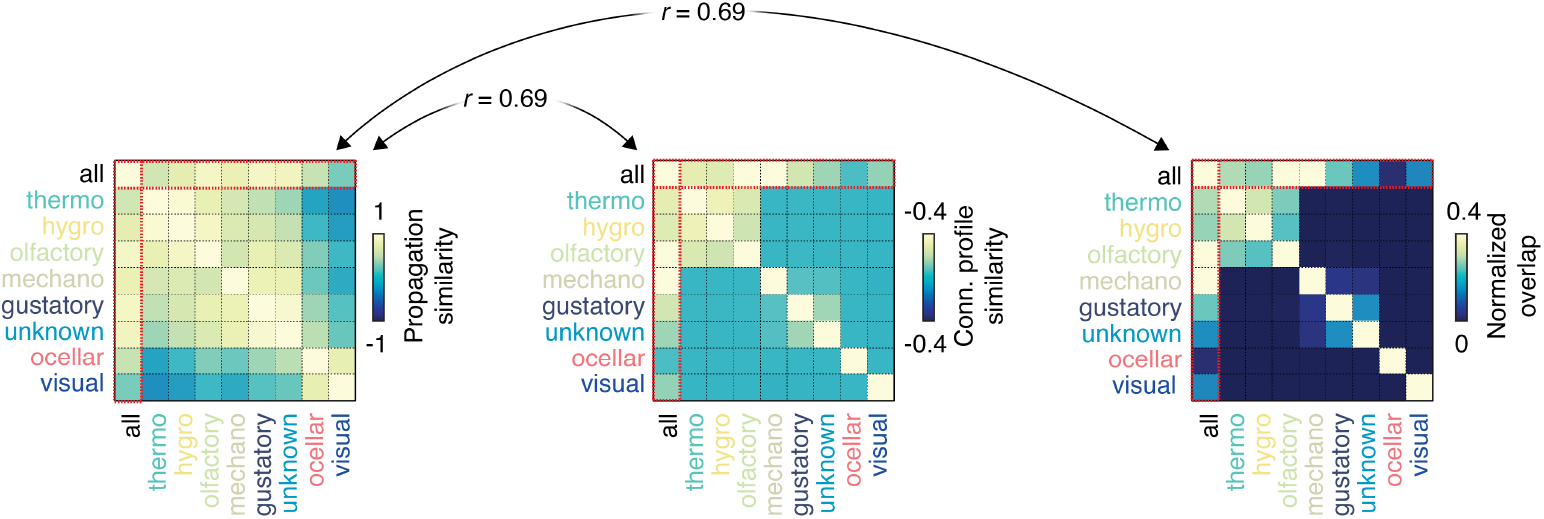
Cascade similarity *versus* connectivity similarity. Here, we show cascade similarity (from Fig. 4) alongside the similarity of connectivity profiles. Specifically, we calculated the an average connectivity profile (outgoing connections only) for all each sensory modalities combined (all) and for each sensory modality independently. The center plot shows correlations between profiles; the right plot shows normalized dot product.

**Figure S4.**
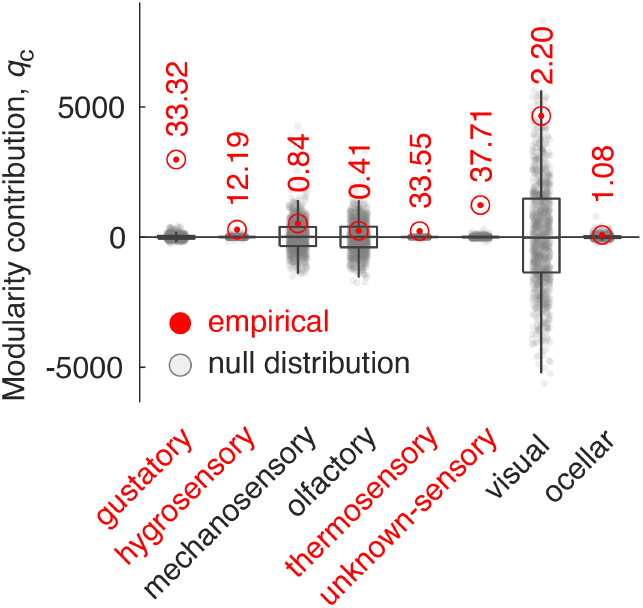
Modularity of sensory systems. We calculated the empirical modularity contribution of each sensory system, *c* as 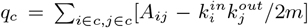. Here *A*_*ij*_ is the number of synapses from neuron *i* to *j*, 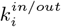 is the weighted in-/out-degree of neuron *i*, and 2*m* = Σ_*i*_ *k*_*i*_ is the total weight of the network. We then compared the observed modularity contribution against a null distribution in which we uniformly and randomly permuted sensory labels (1000 randomizations). We found striking heterogeneity in terms of modularity contributions across modalities; only gustatory, hygrosensory, thermosensory, and unknown sensory neurons were significantly modular (uncorrected *p <* 0.01).

**Figure S5.**
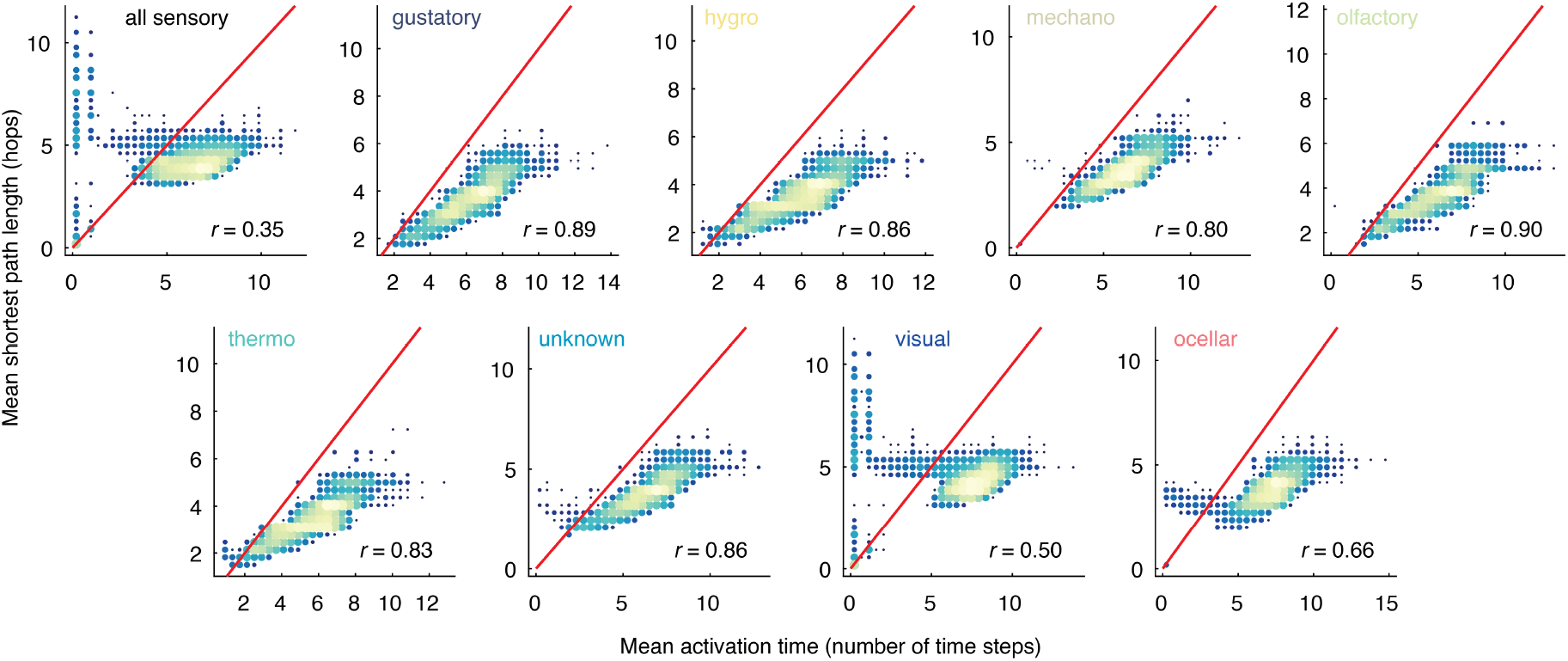
Comparing shortest paths with activation times. In the main text we reported activation times of neurons given different sensory seeds. To verify that these activation times were not strictly the consequence of shortest paths, we calculated the binary shortest path from each neuron to all other neurons in the network. For a given sensory modality, we calculated the average distance of all such sensory neurons to all other neurons in the *Drosophila* brain. We then compared those distances (measured in number of hops) with activation times. The above figures show mean activation times plotted as a function of mean shortest path for all eight sensory modalities as well as all sensory modalities combined. Note that size and color intensity of markers in each plot are proportional to the log density.

**Figure S6.**
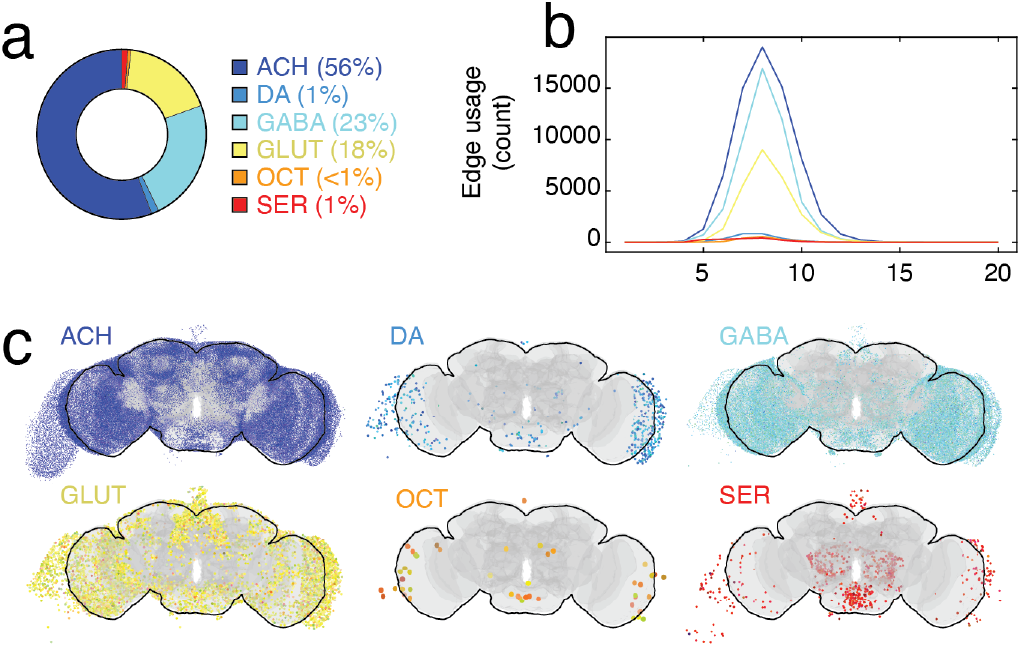
Effect of synapse type on activation time. In the main text we grouped all synapses together, making no distinction between neurotransmitter types. Here, we parse contributions made by six distinct synapse types (based on neurotransmitter). They include acetylcholine (ACH), dopamine (DA), gabaergic (GABA), glutamate (GLUT), octopamine (OCT), and seratonin (SER). (*a*) Percent of all synapses associated with each neurotransmitter. (*b*) We tracked how frequently synapses associated with different neurotransmitter types successfully activated their post-synaptic partner. In this plot, we show edge usage (number of active synapses of each type) as a function of time steps. Note that this plot was generated as the mean usage over 1000 simulations with all sensory modalities included in the seed set but with seed size fixed at 16 with transmission probability of *p*_*tranmission*_ = 0.01. (*c*) For each neuron, we calculated when it was first activated by by what neurotransmitter. This plot depicts, as a function of neurotransmitter, neurons likely to be activated by each of the six neurotransmitters.

**Figure S7.**
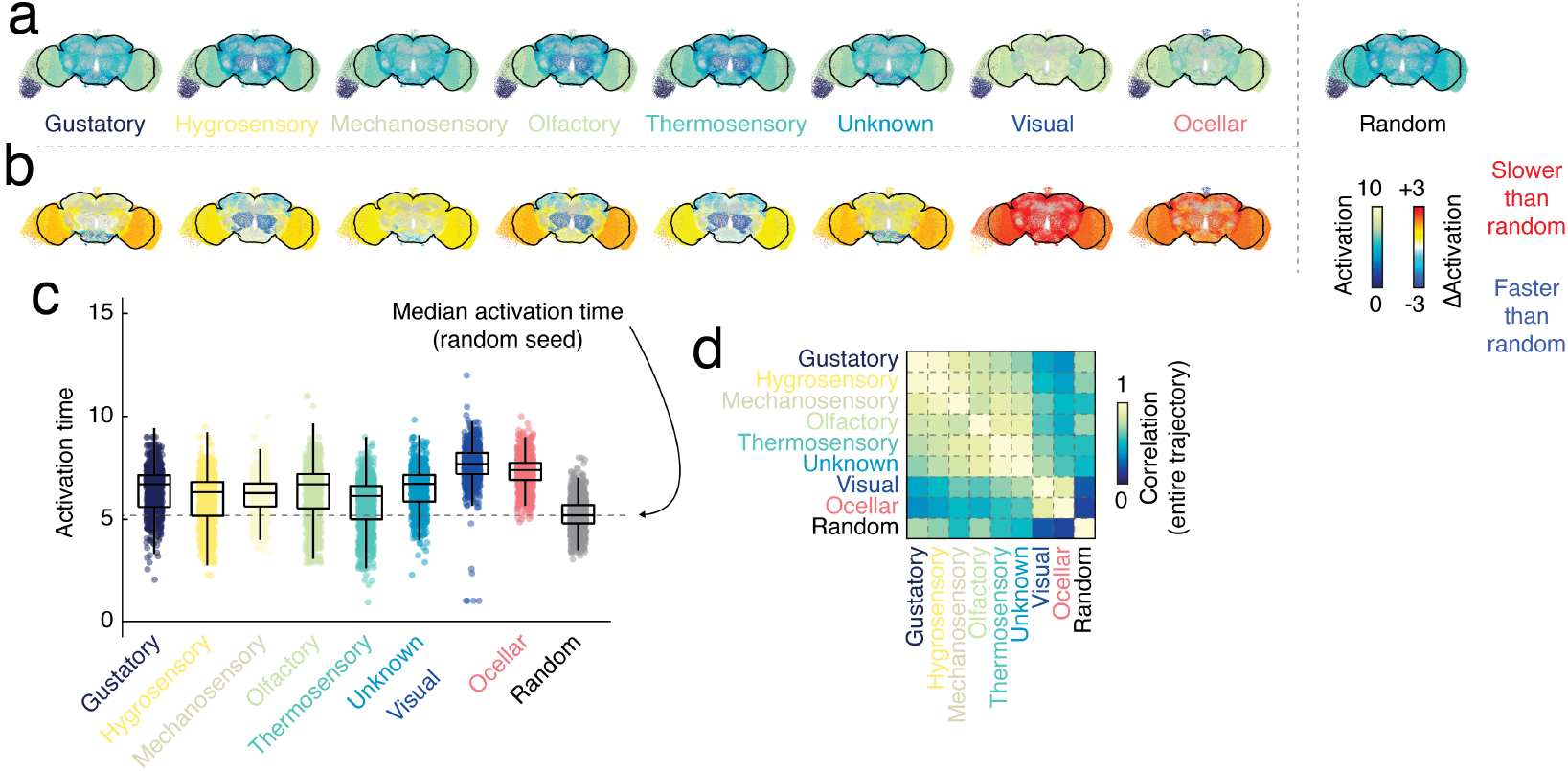
Comparing random and sensory seeds. (*a*) Mean activation time for neurons for eight sensory modalities. To the far right we show activation time for the random map. (*b*) Difference between random and sensory cascades in terms of activation time. (*c*) Boxplot of differences. (*d*) Correlation of entire (vectorized and concatenated) trajectory.

**Figure S8.**
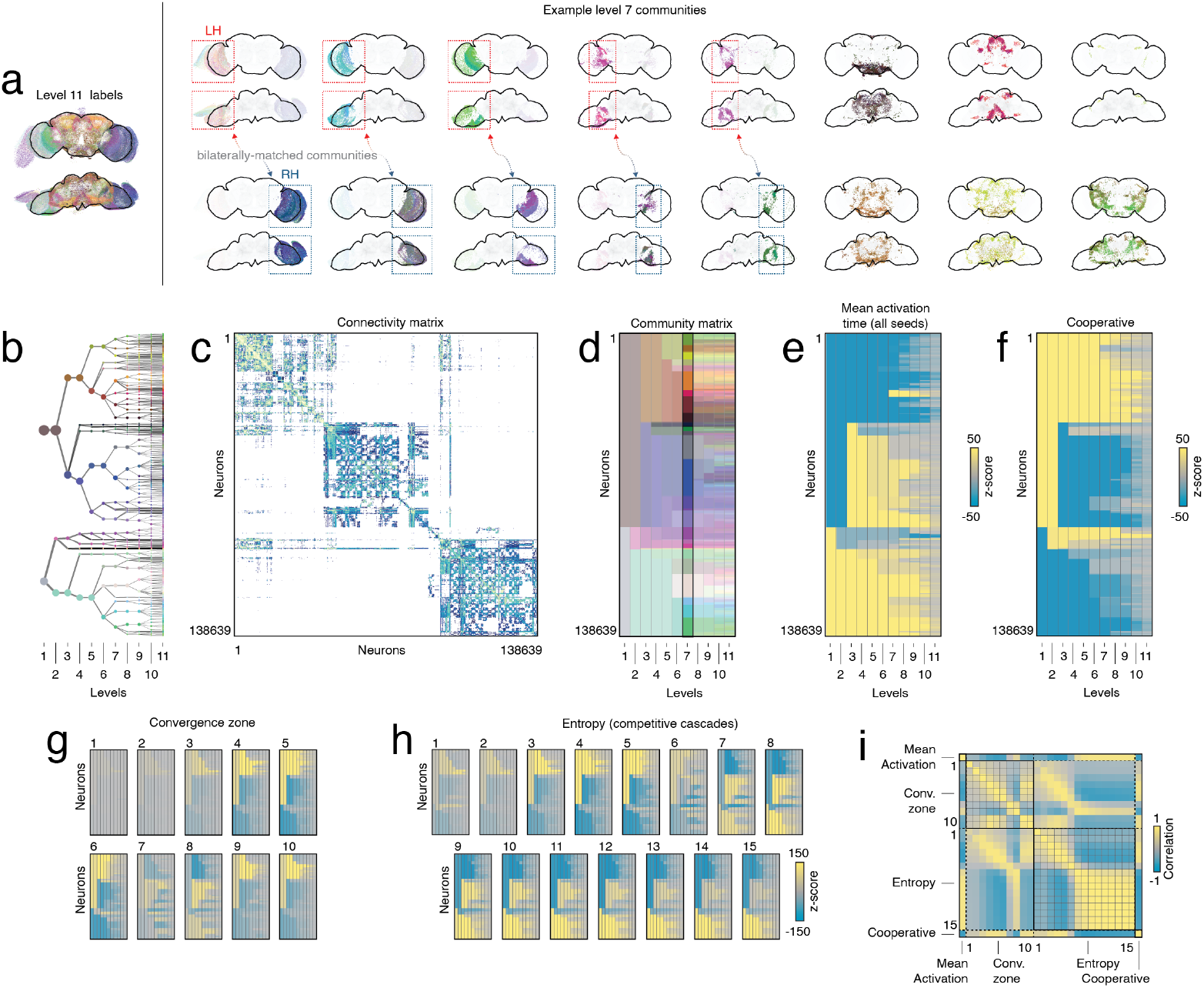
Linking the modular structure of the connectome with cascade statistics. We used a hierarchical community detection algorithm to divide neurons into nested blocks. (*a*) Topography of all communities with their finest (level 11) coloration. To the right, we show example communities from an intermediate scale (level 7). We highlight pairs of communities that are lateralized as well as their partners in the opposite hemisphere. (*b*) Community dendrogram. (*c*) Connectivity matrix ordered by communities. (*d*) An alternative visualization of community labels; each column is hierarchical level with rows ordered by communities. The colors of entries correspond to community labels. Panels *e*-*h* show enrichment scores for communities at all hierarchical levels for: (*e*) Mean activation time; (*f*) Average reduction in activation time during cooperative cascades; (*g*) Convergence zone as a function of time step (from *t* = 1 to *t* = 10), and (*h*) Entropy during competitive cascades (from *t* = 1 to *t* = 15). (*i*) For each measure (and each time step, for those that were temporally resolved), we retained for each community its enrichment score. We represented these scores as vectors and computed the correlation between all such vectors to yield a similarity matrix. The “hot” elements of the matrix identified enrichment patterns that were highly similar; “cooler” elements correspond to dissimilar and anti-correlated enrichment patterns.

**Figure S9.**
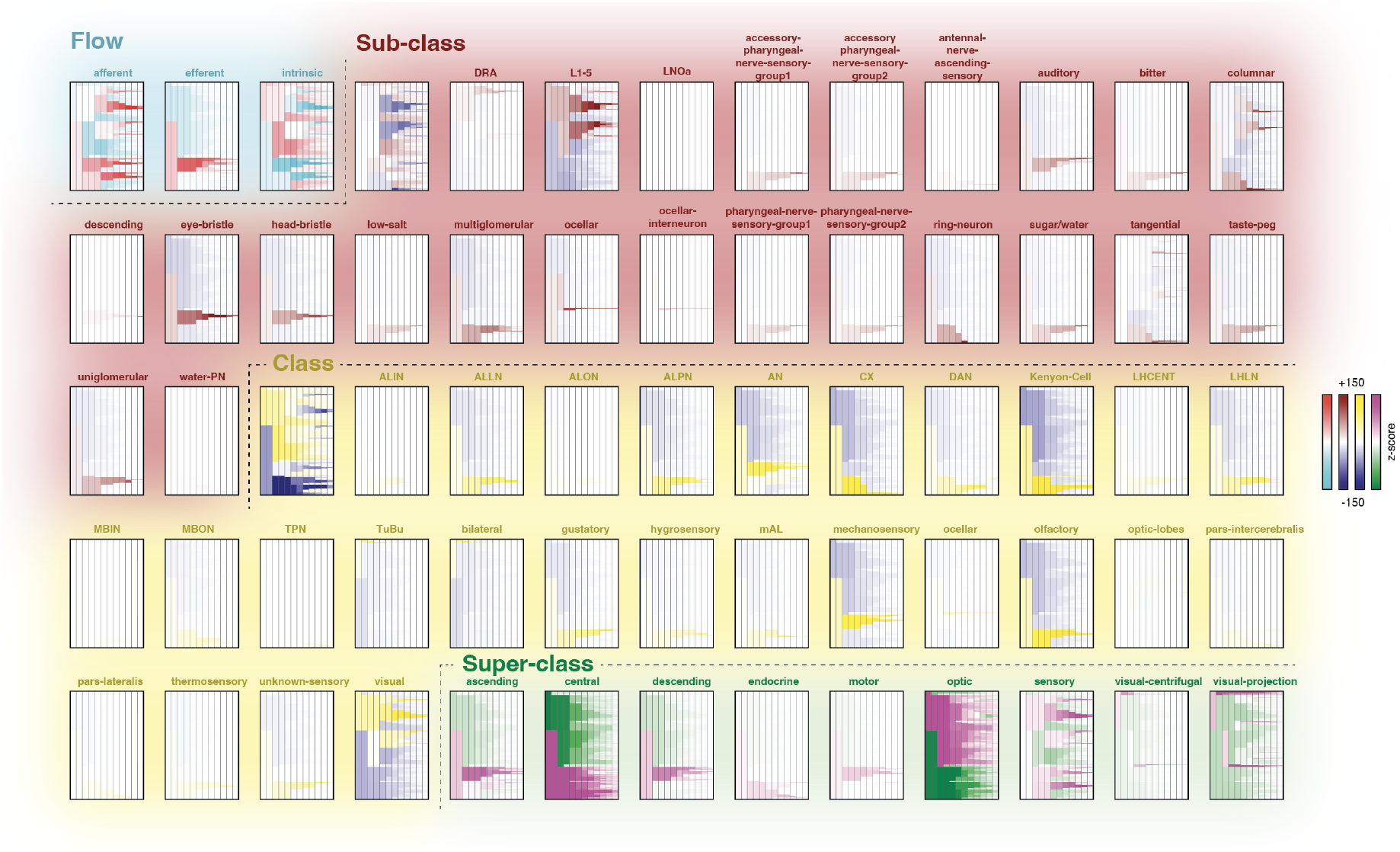
Enrichment analysis of community labels within annotation data. Each plot corresponds to a specific annotation. Each plot shows annotation enrichment across 11 hierarchical levels of communities. The colormap differs across annotations based on annotation category (Flow, sub-class, class, and super-class). Hot colors correspond to communities significantly enriched for an annotation; cooler colors correspond to communities that are avoidant with respect to a given annotation.

**Figure S10.**
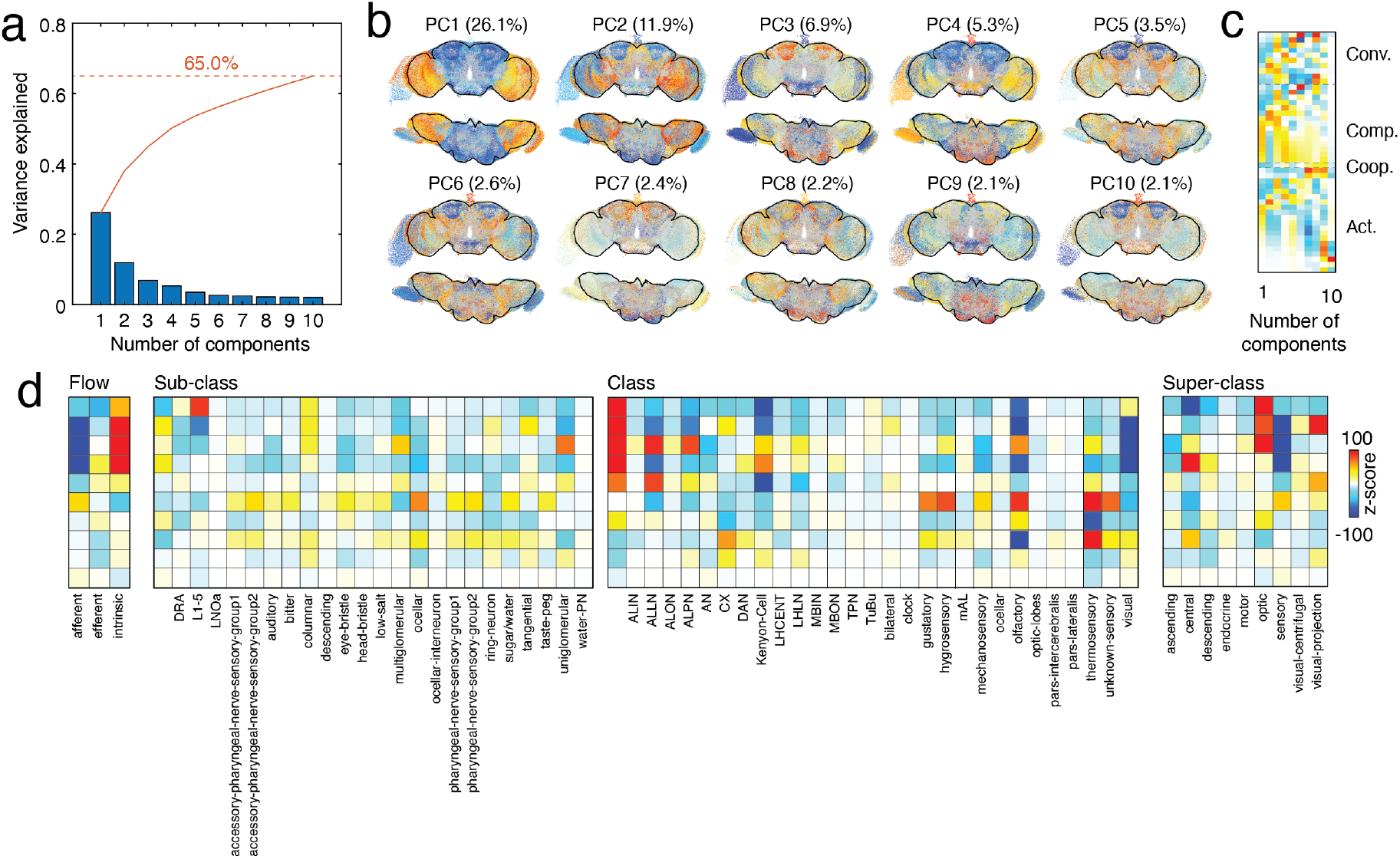
Principal component analysis of cascade statistics. In the main text, we clustered neurons to identify groups with similar cascade statistics–i.e. convergence, speed-ups, entropy, and activation times. Here, we apply PCA to the same data (after z-scoring columns). (*a*) Variance explained by each of the first 10 components (in blue). Cumulative variance explained shown in orange. (*b*) Spatial maps (components). (*c*) Coefficients. (*d*) Enrichment of spatial maps across annotations.

## Notes

### Competing Interest Statement

The authors have declared no competing interest.

### Summary of Updates

Updated figures, added clarifying text, better described the model.

## References

[1] Olaf Sporns, Giulio Tononi, and Rolf Kötter. The human connectome: a structural description of the human brain. PLoS Comput Biol, 1(4):e42, 2005.

[2] Seung Wook Oh, Julie A Harris, Lydia Ng, Brent Winslow, Nicholas Cain, Stefan Mihalas, Quanxin Wang, Chris Lau, Leonard Kuan, Alex M Henry, et al. A mesoscale connectome of the mouse brain. Nature, 508(7495):207–214, 2014.

[3] Larry W Swanson, Joel D Hahn, and Olaf Sporns. Organizing principles for the cerebral cortex network of commissural and association connections. Proceedings of the National Academy of Sciences, 114(45):E9692– E9701, 2017.

[4] Nikola T Markov, MM Ercsey-Ravasz, AR Ribeiro Gomes, Camille Lamy, Loic Magrou, Julien Vezoli, P Misery, A Falchier, R Quilodran, MA Gariel, et al. A weighted and directed interareal connectivity matrix for macaque cerebral cortex. Cerebral cortex, 24(1):17–36, 2014.

[5] Patric Hagmann, Leila Cammoun, Xavier Gigandet, Reto Meuli, Christopher J Honey, Van J Wedeen, and Olaf Sporns. Mapping the structural core of human cerebral cortex. PLoS Biol, 6(7):e159, 2008.

[6] Sven Maximilian Dorkenwald. Analysis of Neuronal Wiring Diagrams. Princeton University, 2023.

[7] Andrea Avena-Koenigsberger, Bratislav Misic, and Olaf Sporns.Communication dynamics in complex brain networks. Nature Reviews Neuroscience, 19(1):17, 2018.

[8] Maxwell H Turner, Kevin Mann, and Thomas R Clandinin. The connectome predicts resting-state functional connectivity across the drosophila brain. Current Biology, 2021.

[9] Francesco Randi, Anuj K Sharma, Sophie Dvali, and Andrew M Leifer. Neural signal propagation atlas of caenorhabditis elegans. Nature, 623(7986):406–414, 2023.

[10] Caio Seguin, Olaf Sporns, and Andrew Zalesky. Brain network communication: concepts, models and applications. Nature Reviews Neuroscience, pages 1–18, 2023.

[11] Cornelia I Bargmann and Eve Marder. From the connectome to brain function. Nature methods, 10(6):483– 490, 2013.

[12] Hae-Jeong Park and Karl Friston. Structural and functional brain networks: from connections to cognition. Science, 342(6158), 2013.

[13] Stéphane Molotchnikoff, Vishal Bharmauria, Lyes Bachatene, Nayan Chanauria, and Jose Fernando Maya-Vetencourt. The function of connectomes in encoding sensory stimuli. Progress in Neurobiology, 181: 101659, 2019.

[14] Nadine Randel, Albina Asadulina, Luis A Bezares-Calderón, Csaba Verasztó, Elizabeth A Williams, Markus Conzelmann, Réza Shahidi, and Gáspár Jékely. Neuronal connectome of a sensory-motor circuit for visual navigation. elife, 3:e02730, 2014.

[15] Danielle S Bassett and Olaf Sporns. Network neuroscience. Nature neuroscience, 20(3):353–364, 2017.

[16] Joshua K Wong, Erik H Middlebrooks, Sanjeet S Grewal, Leonardo Almeida, Christopher W Hess, and Michael S Okun. A comprehensive review of brain connectomics and imaging to improve deep brain stimulation outcomes. Movement Disorders, 35(5):741–751, 2020.

[17] Ed Bullmore and Olaf Sporns. Complex brain networks: graph theoretical analysis of structural and functional systems. Nature reviews neuroscience, 10 (3):186–198, 2009.

[18] Mikail Rubinov and Olaf Sporns. Complex network measures of brain connectivity: uses and interpretations. Neuroimage, 52(3):1059–1069, 2010.

[19] Albert Lin, Runzhe Yang, Sven Dorkenwald, Arie Matsliah, Amy R Sterling, Philipp Schlegel, Szi-chieh Yu, Claire E McKellar, Marta Costa, Katharina Eichler, et al. Network statistics of the whole-brain connectome of drosophila. Nature, 634(8032):153–165, 2024.

[20] Richard F Betzel, John D Medaglia, and Danielle S Bassett. Diversity of meso-scale architecture in human and non-human connectomes. Nature communications, 9(1):1–14, 2018.

[21] Gustavo Deco, Viktor K Jirsa, Peter A Robinson, Michael Breakspear, and Karl Friston. The dynamic brain: from spiking neurons to neural masses and cortical fields. PLoS computational biology, 4(8): e1000092, 2008.

[22] Michael Breakspear. Dynamic models of large-scale brain activity. Nature neuroscience, 20(3):340–352, 2017.

[23] Caio Seguin, Maciej Jedynak, Olivier David, Sina Mansour, Olaf Sporns, and Andrew Zalesky. Communication dynamics in the human connectome shape the cortex-wide propagation of direct electrical stimulation. Neuron, 111(9):1391–1401, 2023.

[24] Janne K Lappalainen, Fabian D Tschopp, Sridhama Prakhya, Mason McGill, Aljoscha Nern, Kazunori Shinomiya, Shin-ya Takemura, Eyal Gruntman, Jakob H Macke, and Srinivas C Turaga. Connectome-constrained networks predict neural activity across the fly visual system. Nature, pages 1–9, 2024.

[25] Duncan J Watts. A simple model of global cascades on random networks. Proceedings of the National Academy of Sciences, 99(9):5766–5771, 2002.

[26] Mark Granovetter. Threshold models of collective behavior. American journal of sociology, 83(6):1420– 1443, 1978.

[27] Bratislav Mišić, Richard F Betzel, Azadeh Nematzadeh, Joaquin Goni, Alessandra Griffa, Patric Hagmann, Alessandro Flammini, Yong-Yeol Ahn, and Olaf Sporns. Cooperative and competitive spreading dynamics on the human connectome. Neuron, 86(6): 1518–1529, 2015.

[28] Kamal Shadi, Eva Dyer, and Constantine Dovrolis. Multisensory integration in the mouse cortical connectome using a network diffusion model. Network Neuroscience, 4(4):1030–1054, 2020.

[29] Jacob C Worrell, Jeffrey Rumschlag, Richard F Betzel, Olaf Sporns, and Bratislav Mišić. Optimized connectome architecture for sensory-motor integration. Network Neuroscience, 1(4):415–430, 2017.

[30] John G White, Eileen Southgate, J Nichol Thomson, and Sydney Brenner. The structure of the nervous system of the nematode caenorhabditis elegans. Philos Trans R Soc Lond B Biol Sci, 314(1165):1–340, 1986.

[31] Moritz Helmstaedter, Kevin L Briggman, Srinivas C Turaga, Viren Jain, H Sebastian Seung, and Winfried Denk. Connectomic reconstruction of the inner plexiform layer in the mouse retina. Nature, 500(7461): 168–174, 2013.

[32] Ann-Shyn Chiang, Chih-Yung Lin, Chao-Chun Chuang, Hsiu-Ming Chang, Chang-Huain Hsieh, Chang-Wei Yeh, Chi-Tin Shih, Jian-Jheng Wu, Guo-Tzau Wang, Yung-Chang Chen, et al. Threedimensional reconstruction of brain-wide wiring networks in drosophila at single-cell resolution. Current biology, 21(1):1–11, 2011.

[33] Alexander Shapson-Coe, Michał Januszewski Daniel R Berger, Art Pope, Yuelong Wu, Tim Blakely, Richard L Schalek, Peter H Li, Shuohong Wang, Jeremy Maitin-Shepard, et al. A petavoxel fragment of human cerebral cortex reconstructed at nanoscale resolution. Science, 384(6696):eadk4858, 2024.

[34] Mojtaba R Tavakoli, Julia Lyudchik, Michał Januszewski Vitali Vistunou, Nathalie Agudelo, Jakob Vorlaufer, Christoph Sommer, Caroline Kreuzinger, Barbara Oliveira, Alban Cenameri, et al. Light-microscopy based dense connectomic reconstruction of mammalian brain tissue. bioRxiv, pages 2024–03, 2024.

[35] Sven Dorkenwald, Arie Matsliah, Amy R Sterling, Philipp Schlegel, Szi-Chieh Yu, Claire E McKellar, Albert Lin, Marta Costa, Katharina Eichler, Yijie Yin, et al. Neuronal wiring diagram of an adult brain. Nature, 634(8032):124–138, 2024.

[36] Louis K Scheffer, C Shan Xu, Michal Januszewski, Zhiyuan Lu, Shin-ya Takemura, Kenneth J Hayworth, Gary B Huang, Kazunori Shinomiya, Jeremy Maitlin-Shepard, Stuart Berg, et al. A connectome and analysis of the adult drosophila central brain. Elife, 9: e57443, 2020.

[37] Philipp Schlegel, Yijie Yin, Alexander S Bates, Sven Dorkenwald, Katharina Eichler, Paul Brooks, Daniel S Han, Marina Gkantia, Marcia Dos Santos, Eva J Munnelly, et al. Whole-brain annotation and multi-connectome cell typing of drosophila. Nature, 634 (8032):139–152, 2024.

[38] Nicole K Guittari, Miguel E Wimbish, Patricia K Rivlin, Mark A Hinton, Jordan K Matelsky, Victoria A Rose, Jorge L Rivera Jr, Nicole E Stock, Brock A Wester, Erik C Johnson, et al. Nanoscale connectomics annotation standards framework. arXiv preprint 2410.22320, 2024.

[39] Katharina Eichler, Feng Li, Ashok Litwin-Kumar, Youngser Park, Ingrid Andrade, Casey M Schneider-Mizell, Timo Saumweber, Annina Huser, Claire Eschbach, Bertram Gerber, et al. The complete connectome of a learning and memory centre in an insect brain. Nature, 548(7666):175–182, 2017.

[40] Shahar Frechter, Alexander Shakeel Bates, Sina Tootoonian, Michael-John Dolan, James Manton, Arian Rokkum Jamasb, Johannes Kohl, Davi Bock, and Gregory Jefferis. Functional and anatomical specificity in a higher olfactory centre. elife, 8:e44590, 2019.

[41] C Shan Xu, Michal Januszewski, Zhiyuan Lu, Shinya Takemura, Kenneth J Hayworth, Gary Huang, Kazunori Shinomiya, Jeremy Maitin-Shepard, David Ackerman, Stuart Berg, et al. A connectome of the adult drosophila central brain. BioRxiv, pages 2020– 01, 2020.

[42] Yi Sun, Aljoscha Nern, Romain Franconville, Hod Dana, Eric R Schreiter, Loren L Looger, Karel Svoboda, Douglas S Kim, Ann M Hermundstad, and Vivek Jayaraman. Neural signatures of dynamic stimulus selection in drosophila. Nature neuroscience, 20 (8):1104–1113, 2017.

[43] Michael Winding, Benjamin D Pedigo, Christopher L Barnes, Heather G Patsolic, Youngser Park, Tom Kazimiers, Akira Fushiki, Ingrid V Andrade, Avinash Khandelwal, Javier Valdes-Aleman, et al. The connectome of an insect brain. Science, 379(6636):eadd9330, 2023.

[44] Olaf Sporns. Structure and function of complex brain networks. Dialogues in clinical neuroscience, 15(3): 247–262, 2013.

[45] Azadeh Nematzadeh, Emilio Ferrara, Alessandro Flammini, and Yong-Yeol Ahn. Optimal network modularity for information diffusion. Physical review letters, 113(8):088701, 2014.

[46] Raphael Cohn, Ianessa Morantte, and Vanessa Ruta. Coordinated and compartmentalized neuromodulation shapes sensory processing in drosophila. Cell, 163(7): 1742–1755, 2015.

[47] Tanya Wolff and Gerald M Rubin. Neuroarchitecture of the drosophila central complex: A catalog of nodulus and asymmetrical body neurons and a revision of the protocerebral bridge catalog. Journal of Comparative Neurology, 526(16):2585–2611, 2018.

[48] Brad K Hulse, Hannah Haberkern, Romain Franconville, Daniel B Turner-Evans, Shin-ya Takemura, Tanya Wolff, Marcella Noorman, Marisa Dreher, Chuntao Dan, Ruchi Parekh, et al. A connectome of the drosophila central complex reveals network motifs suitable for flexible navigation and context-dependent action selection. Elife, 10, 2021.

[49] Dominic D Frank, Anders Enjin, Genevieve C Jouandet, Emanuela E Zaharieva, Alessia Para, Marcus C Stensmyr, and Marco Gallio. Early integration of temperature and humidity stimuli in the drosophila brain. Current Biology, 27(15):2381–2388, 2017.

[50] Edsger W Dijkstra. A note on two problems in connexion with graphs. In Edsger Wybe Dijkstra: his life, work, and legacy, pages 287–290. 2022.

[51] Tiago P Peixoto. Hierarchical block structures and high-resolution model selection in large networks. Physical Review X, 4(1):011047, 2014.

[52] Richard Betzel, Maria Grazia Puxeddu, and Caio Seguin. Hierarchical communities in the larval drosophila connectome: Links to cellular annotations and network topology. Proceedings of the National Academy of Sciences, 121(38):e2320177121, 2024.

[53] Ketan Mehta, Rebecca F Goldin, and Giorgio A Ascoli. Circuit analysis of the drosophila brain using connectivity-based neuronal classification reveals organization of key communication pathways. Network Neuroscience, 7(1):269–298, 2023.

[54] Alexander B Kunin, Jiahao Guo, Kevin E Bassler, Xaq Pitkow, and Krešimir Josić. Hierarchical modular structure of the drosophila connectome. Journal of Neuroscience, 43(37):6384–6400, 2023.

[55] Leandro González-Montesino, Darian H Grass-Boada, and Rubén Armañnazas. Network community detection in connectomics data using graph theory. In 2023 IEEE International Conference on Bioinformatics and Biomedicine (BIBM), pages 3459–3465. IEEE, 2023.

[56] Hee-Sun J. Cheong et al. Multi-regional circuits underlying visually guided decision making in Drosophila. Neuron, 106(4):602–616, 2020. doi: 10.1016/j.neuron.2020.02.020.

[57] Dean A. Pospisil, Max J. Aragon, Sven Dorkenwald, et al. The fly connectome reveals a path to the effectome. Nature, 634:201–207, 2024. doi:10.1038/s41586-024-07982-0.

[58] Albert Lin, Runzhe Yang, Sven Dorkenwald, et al. Network statistics of the whole-brain connectome of drosophila. Nature, 634:186–193, 2024. doi: 10.1038/s41586-024-07968-y.

[59] Philip K Shiu, Gabriella R Sterne, Nico Spiller, Romain Franconville, Andrea Sandoval, Joie Zhou, Neha Simha, Chan Hyuk Kang, Seongbong Yu, Jinseop S Kim, et al. A drosophila computational brain model reveals sensorimotor processing. Nature, 634(8032): 210–219, 2024.

[60] Apostolos Fotiadis et al. Building and integrating brain-wide maps of nervous system function. Science, 376:eabm5000, 2024. doi:10.1126/science.abm5000.

[61] Bryan O. Turner et al. The connectome and beyond: structural–functional coupling in the brain. Nature Reviews Neuroscience, 22:1–16, 2021. doi: 10.1038/s41583-021-00466-4.

[62] Teemu Lappalainen et al. Learning synaptic weights in the fly connectome: implications for structure–function mapping. bioRxiv, 2024. doi: 10.1101/2024.05.20.601123.

[63] Lyle Muller, Frédéric Chavane, John Reynolds, and Terrence J Sejnowski. Cortical travelling waves: mechanisms and computational principles. Nature Reviews Neuroscience, 19(5):255–268, 2018.

[64] Selen Atasoy, Isaac Donnelly, and Joel Pearson. Human brain networks function in connectome-specific harmonic waves. Nature communications, 7(1):10340, 2016.

[65] Romualdo Pastor-Satorras, Claudio Castellano, Piet Van Mieghem, and Alessandro Vespignani. Epidemic processes in complex networks. Reviews of modern physics, 87(3):925–979, 2015.

[66] Marcus Ghosh, Gabriel Béna, Volker Bormuth, and Dan FM Goodman. Nonlinear fusion is optimal for a wide class of multisensory tasks. PLOS Computational Biology, 20(7):e1012246, 2024.

[67] Sibo Wang-Chen, Victor Alfred Stimpfling, Thomas Ka Chung Lam, Pembe Gizem Özdil, Louise Genoud, Femke Hurtak, and Pavan Ramdya. Neuromechfly v2: simulating embodied sensorimotor control in adult drosophila. Nature Methods, 21(12):2353–2362, 2024.

[68] Roman Vaxenburg, Igor Siwanowicz, Josh Merel, Alice A Robie, Carmen Morrow, Guido Novati, Zinovia Stefanidi, Gert-Jan Both, Gwyneth M Card, Michael B Reiser, et al. Whole-body physics simulation of fruit fly locomotion. Nature, pages 1–3, 2025.

[69] Gorka Zamora-López and Matthieu Gilson. An integrative dynamical perspective for graph theory and the analysis of complex networks. Chaos: An Interdisciplinary Journal of Nonlinear Science, 34(4), 2024.

[70] Zhihao Zheng, J Scott Lauritzen, Eric Perlman, Camenzind G Robinson, Matthew Nichols, Daniel Milkie, Omar Torrens, John Price, Corey B Fisher, Nadiya Sharifi, et al. A complete electron microscopy volume of the brain of adult drosophila melanogaster. Cell, 174(3):730–743, 2018.

[71] Sven Dorkenwald, Claire E McKellar, Thomas Macrina, Nico Kemnitz, Kisuk Lee, Ran Lu, Jingpeng Wu, Sergiy Popovych, Eric Mitchell, Barak Nehoran, et al. Flywire: online community for whole-brain connectomics. Nature methods, 19(1):119–128, 2022.

[72] Julia Buhmann, Arlo Sheridan, Caroline Malin-Mayor, Philipp Schlegel, Stephan Gerhard, Tom Kazimiers, Renate Krause, Tri M Nguyen, Larissa Heinrich, WeiChung Allen Lee, et al. Automatic detection of synaptic partners in a whole-brain drosophila electron microscopy data set. Nature methods, 18(7):771–774, 2021.

[73] Larissa Heinrich, Jan Funke, Constantin Pape, Juan Nunez-Iglesias, and Stephan Saalfeld. Synaptic cleft segmentation in non-isotropic volume electron microscopy of the complete drosophila brain. In Medical Image Computing and Computer Assisted Intervention–MICCAI 2018: 21st International Conference, Granada, Spain, September 16-20, 2018, Proceedings, Part II 11, pages 317–325. Springer, 2018.

[74] Arie Matsliah, Szi-Chieh Yu, Krzysztof Kruk, Doug Bland, Austin T Burke, Jay Gager, James Hebditch, Ben Silverman, Kyle Patrick Willie, Ryan Willie, et al. Neuronal parts list and wiring diagram for a visual system. Nature, 634(8032):166–180, 2024.

[75] Taro Kaneuchi, Caroline V Sartain, Satomi Takeo, Vanessa L Horner, Norene A Buehner, Toshiro Aigaki, and Mariana F Wolfner. Calcium waves occur as drosophila oocytes activate. Proceedings of the National Academy of Sciences, 112(3):791–796, 2015.

[76] Ben Jiwon Choi, Yen-Chung Chen, and Claude Desplan. Retinal calcium waves coordinate uniform tissue patterning of the drosophila eye. Science, 390(6775): eady5541, 2025.

[77] Takaaki Matsui. Calcium wave propagation during cell extrusion. Current Opinion in Cell Biology, 76:102083, 2022.

[78] Tiago Branco and Kevin Staras. The probability of neurotransmitter release: variability and feedback control at single synapses. Nature Reviews Neuroscience, 10(5):373–383, 2009.

[79] Venkatesh N Murthy, Thomas Schikorski, Charles F Stevens, and Yi Zhu. Inactivity produces increases in neurotransmitter release probability and synapse size. Neuron, 18(4):599–612, 1997.

[80] J Del Castillo and B Katz. Quantal components of the end-plate potential. The Journal of Physiology, 124(3):560–573, 1954.

[81] Thomas Schikorski and Charles F Stevens. Quantitative ultrastructural analysis of hippocampal excitatory synapses. The Journal of Neuroscience, 17(15):5858– 5867, 1997.

[82] Shin Ya Takemura, Arpit Bharioke, Zheng Lu, Aljoscha Nern, Snigdha Vitaladevuni, Patricia K Rivlin, Winston T Katz, Donald J Olbris, Stephen Plaza, Philip Winston, et al. A visual motion detection circuit suggested by drosophila connectomics. Nature, 500(7461):175–181, 2013.

[83] Jeffrey C Magee. Dendritic integration of excitatory synaptic input. Nature Reviews Neuroscience, 1(3): 181–190, 2000.

[84] Jeffry S. Isaacson and Massimo Scanziani. How inhibition shapes cortical activity. Neuron, 72(2):231–243, 2011.

[85] L.F. Abbott, J.A. Varela, K. Sen, and S.B. Nelson. Synaptic depression and cortical gain control. Science, 275(5297):220–224, 1997.

[86] Amos Arieli, Anna Sterkin, Amiram Grinvald, and Ad Aertsen. Dynamics of ongoing activity: explanation of the large variability in evoked cortical responses. Science, 273(5283):1868–1871, 1996.

[87] Vikas Bhandawat, Shawn R Olsen, Michele L Schlief, Nathaniel W Gouwens, and Rachel I Wilson. Sensory processing in the drosophila antennal lobe increases reliability and separability of ensemble odor representations. Nature Neuroscience, 10:1474–1482, 2007.

[88] Rachel I Wilson, Glenn C Turner, and Gilles Laurent. Transformation of olfactory representations in the drosophila antennal lobe. Neuron, 44(2):303–314, 2004.

[89] Bruno A Olshausen and David J Field. Sparse coding of sensory inputs. Current Opinion in Neurobiology, 14(4):481–487, 2004.

[90] David H Hubel and Torsten N Wiesel. Receptive fields and functional architecture of monkey striate cortex. The Journal of Physiology, 195(1):215–243, 1968.

[91] Gwyneth M Card and Michael H Dickinson. Visually mediated motor planning in the escape response of drosophila. Current Biology, 18(17):1300–1307, 2008.

[92] Ian Nauhaus, Laura Busse, Dario L Ringach, and Matteo Carandini. Stimulus contrast modulates functional connectivity in visual cortex. Nature Neuroscience, 12: 70–76, 2009.

[93] György Buzsáki. Hippocampal sharp wave–ripple: A cognitive biomarker for episodic memory and planning. Hippocampus, 25(10):1073–1188, 2015.

[94] Wei-Chung Allen Lee, Vincent Bonin, Michael Reed, Brett J Graham, Greg Hood, Katie Glattfelder, and R Clay Reid. Anatomy and function of an excitatory network in the visual cortex. Nature, 532(7599):370– 374, 2016.

[95] Jonathan D Power, Alexander L Cohen, Steven M Nelson, Gagan S Wig, Kelly Anne Barnes, Jessica A Church, Alecia C Vogel, Timothy O Laumann, Fran M Miezin, Bradley L Schlaggar, et al. Functional network organization of the human brain. Neuron, 72(4):665– 678, 2011.

[96] David Meunier, Renaud Lambiotte, and Edward T Bullmore. Modular and hierarchically modular organization of brain networks. Frontiers in neuroscience, 4:200, 2010.

[97] Richard F Betzel, Katherine C Wood, Christopher Angeloni, Maria Neimark Geffen, and Danielle S Bassett. Stability of spontaneous, correlated activity in mouse auditory cortex. PLoS computational biology, 15(12): e1007360, 2019.

[98] Richard F Betzel. Organizing principles of whole-brain functional connectivity in zebrafish larvae. Network Neuroscience, 4(1):234–256, 2020.

[99] Travis A Jarrell, Yi Wang, Adam E Bloniarz, Christopher A Brittin, Meng Xu, J Nichol Thomson, Donna G Albertson, David H Hall, and Scott W Emmons. The connectome of a decision-making neural network. science, 337(6093):437–444, 2012.

[100] Dragana M Pavlovic, Petra E Vértes, Edward T Bullmore, William R Schafer, and Thomas E Nichols. Stochastic blockmodeling of the modules and core of the caenorhabditis elegans connectome. PloS one, 9 (7):e97584, 2014.

[101] Daiki Kiyooka, Ikumi Oomoto, Jun Kitazono, Midori Kobayashi, Chie Matsubara, Kenta Kobayashi, Masanori Murayama, and Masafumi Oizumi. Singlecell resolution functional networks during sleep are segregated into spatially intermixed modules. bioRxiv, pages 2023–09, 2023.

[102] Santo Fortunato. Community detection in graphs. Physics reports, 486(3-5):75–174, 2010.

[103] Mark EJ Newman and Michelle Girvan. Finding and evaluating community structure in networks. Physical review E, 69(2):026113, 2004.

[104] Martin Rosvall and Carl T Bergstrom. Maps of random walks on complex networks reveal community structure. Proceedings of the National Academy of Sciences, 105(4):1118–1123, 2008.

[105] Pan Zhang and Cristopher Moore. Scalable detection of statistically significant communities and hierarchies, using message passing for modularity. Proceedings of the National Academy of Sciences, 111(51): 18144–18149, 2014.

[106] Gergely Palla, Imre Derényi, Illés Farkas, and Tamás Vicsek. Uncovering the overlapping community structure of complex networks in nature and society. nature, 435(7043):814–818, 2005.

[107] Sadamori Kojaku, Filippo Radicchi, Yong-Yeol Ahn, and Santo Fortunato. Network community detection via neural embeddings. Nature Communications, 15 (1):9446, 2024.

[108] Carey E Priebe, Youngser Park, Joshua T Vogelstein, John M Conroy, Vince Lyzinski, Minh Tang, Avanti Athreya, Joshua Cape, and Eric Bridgeford. On a two-truths phenomenon in spectral graph clustering. Proceedings of the National Academy of Sciences, 116 (13):5995–6000, 2019.

[109] Brian Karrer and Mark EJ Newman. Stochastic block-models and community structure in networks. Physical review E, 83(1):016107, 2011.

[110] Paul W Holland, Kathryn Blackmond Laskey, and Samuel Leinhardt. Stochastic blockmodels: First steps. Social networks, 5(2):109–137, 1983.

[111] Carolyn J Anderson, Stanley Wasserman, and Katherine Faust. Building stochastic blockmodels. Social networks, 14(1-2):137–161, 1992.

[112] Tom AB Snijders and Krzysztof Nowicki. Estimation and prediction for stochastic blockmodels for graphs with latent block structure. Journal of classification, 14(1):75–100, 1997.

[113] Krzysztof Nowicki and Tom A B Snijders. Estimation and prediction for stochastic blockstructures. Journal of the American statistical association, 96(455):1077– 1087, 2001.

[114] Leto Peel, Daniel B Larremore, and Aaron Clauset. The ground truth about metadata and community detection in networks. Science advances, 3(5):e1602548, 2017.

[115] Richard F Betzel, Maxwell A Bertolero, and Danielle S Bassett. Non-assortative community structure in resting and task-evoked functional brain networks. bioRxiv, page 355016, 2018.

[116] Joshua Faskowitz, Xiaoran Yan, Xi-Nian Zuo, and Olaf Sporns. Weighted stochastic block models of the human connectome across the life span. Scientific reports, 8(1):1–16, 2018.

[117] Christopher Aicher, Abigail Z Jacobs, and Aaron Clauset. Adapting the stochastic block model to edge-weighted networks. arXiv preprint 1305.5782, 2013.

[118] Marcus E Raichle. The brain’s default mode network. Annual review of neuroscience, 38:433–447, 2015.

[119] Tiago P Peixoto. Entropy of stochastic blockmodel ensembles. Physical Review E, 85(5):056122, 2012.

[120] Tiago P Peixoto. Efficient monte carlo and greedy heuristic for the inference of stochastic block models. Physical Review E, 89(1):012804, 2014.

[121] Tiago P Peixoto. Merge-split markov chain monte carlo for community detection. Physical Review E, 102 (1):012305, 2020.

[122] Andrea Lancichinetti and Santo Fortunato. Consensus clustering in complex networks. Scientific reports, 2 (1):1–7, 2012.

[123] Tiago P Peixoto. Revealing consensus and dissensus between network partitions. Physical Review X, 11(2): 021003, 2021.

[124] J MacQueen. Some methods for classification and analysis of multivariate observations. In Proceedings of 5-th Berkeley Symposium on Mathematical Statistics and Probability/University of California Press, 1967.

